## Supplementary information for "Ten-eleven translocation 1 Mediated-DNA Hydroxymethylation is Required for Myelination and Remyelination in the Mouse Brain"

**Supplementary figures:**

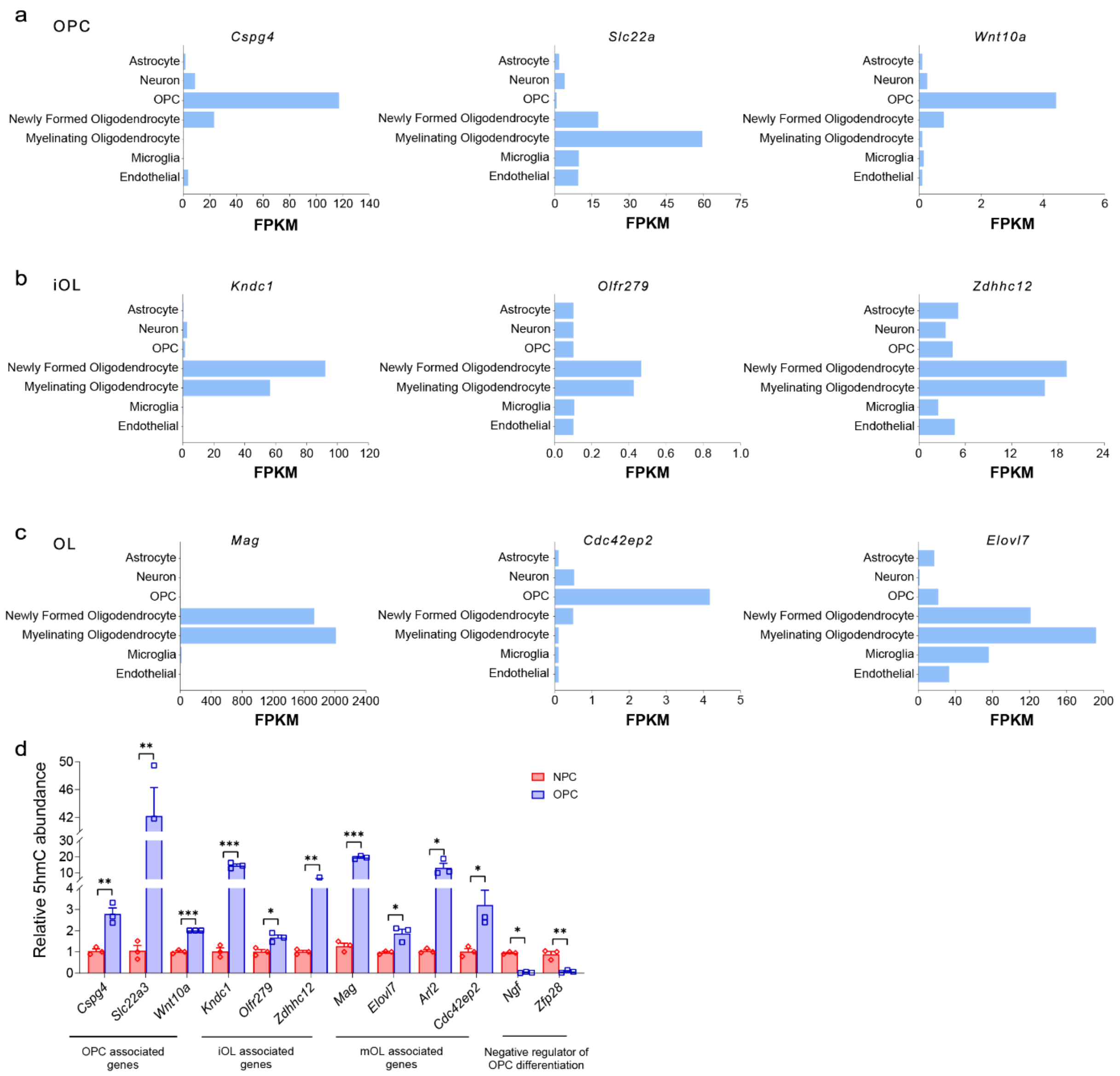


**Supplementary Fig. 1 Distribution pattern of genomic 5hmCs is associated with the transition from NPCs to OPCs**

(a-c) Examples of OL lineage-specific genes expressed in different OL stages. Fragments per kilobase of transcript per million mapped reads (FPKMs) of representative genes highly expressed in OPCs (a), iOLs (b), and OLs (c). Diagrams were made from http://jiaqianwulab.org/braincell/RNASeq.html.

(d) The level of 5hmC in certain gene loci is identified by qPCR assay. Validation of loci specific 5hmC modification in OPC and NPC cultures by qPCR analysis of OPC associated genes, immature OLs (iOL) associated genes, mOL associated genes and negative regulators of OPC differentiation. Data are Means ± SEM ($n$ = 3 independent cultures per group). *, $p$<0.05, **, $p$<0.01, ***, $p$<0.001, compared to NPC. Two-tailed unpaired $t$ test. *Cspg4*: $t$ = -5.867, $df$ = 4, $p$ = 0.004; *Slc22a3*: $t$ = -10.100, $df$ = 4, $p$ = 0.001; *Wnt10a*: $t$ = -20.501, $df$ = 4, $p$ = 0.000033; *Kndc1*: $t$ = -13.045, $df$ = 4, $p$ = 0.000199; *Olfr279*: $t$ = -4.221, $df$ = 4, $p$ = 0.013; *Zdhhc12*: $t$ = -5.173, $df$ = 4, $p$ = 0.007; *Mag*: $t$ = -32.333, $df$ = 4, $p$ = 0.000005; *Elovl7*: $t$ = -4.274, $df$ = 4, $p$ = 0.013; *Arl2*: $t$ = -4.319, $df$ = 4, $p$ = 0.012; *Cdc42ep2*: $t$ = -3.081, $df$ = 4, $p$ = 0.037; *Ngf*: $t$ = 27.294, $df$ = 4, $p$ = 0.037; *Zfp28*: $t$ = 5.877, $df$ = 4, $p$ = 0.004.

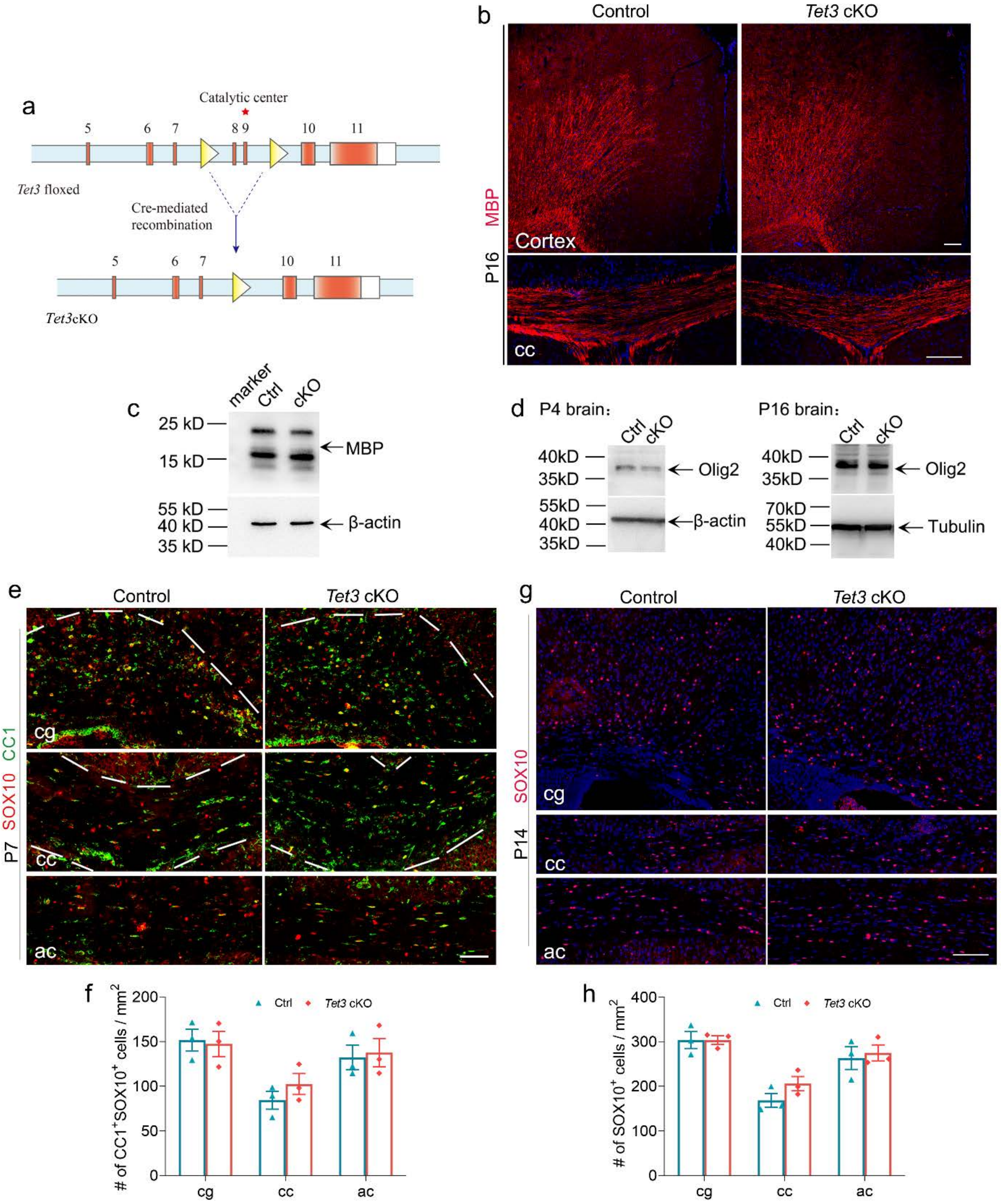


**Supplementary Fig. 2 Deletion of *Tet3* does not cause significant effects on myelination**

(a) Schematic diagram of Cre-mediated excision of floxed *Tet3* exons encoding the catalytic center.

(b) Representative image of MBP immunostaining of postnatal day 16 (P16) brains from control and *Tet3* cKO mice. Scale bar, 100 μm.

(c) Western blot assay for MBP expression in P16 control (Ctrl) and *Tet3* cKO (cKO) brains. β-actin was detected as loading control.

(d) Western blot assay for OLIG2 expression in P4 and P16 control and *Tet3* cKO brains. β-actin and Tubulin was detected as loading control, respectively.

(e) Representative images of three regions of P7 control and *Tet3* cKO brains stained for SOX10 and CC1. Scale bar, 100 μm. cc; corpus callosum; cg, cingulum; ac, anterior commissure.

(f) Density of SOX10$^+$CC1$^+$ cells in P7 control and *Tet3* cKO brains. Data are Means ± SEM (*n* =

3 animals per group). Two-tailed unpaired *t* test, cg: $t = 0.226$, $df = 4$, $p = 0.833$; cc: $t = -0.170$, $df = 4$, $p = 0.307$; ac: $t = -0.257$, $df = 4$, $p = 0.810$.
(g) Representative images of three regions of P14 control and *Tet3* cKO brains stained for SOX10. Scale bar, 100 μm.
(h) Quantification of $SOX10^+$ cells in P14 control and *Tet3* cKO brains. Data are Means ± SEM ($n = 3$ animals per group). Two-tailed unpaired *t* test, cg: $t = 0.012$, $df = 4$, $p = 0.991$; cc: $t = -1.684$, $df = 4$, $p = 0.168$; ac: $t = -0.380$, $df = 4$, $p = 0.723$.

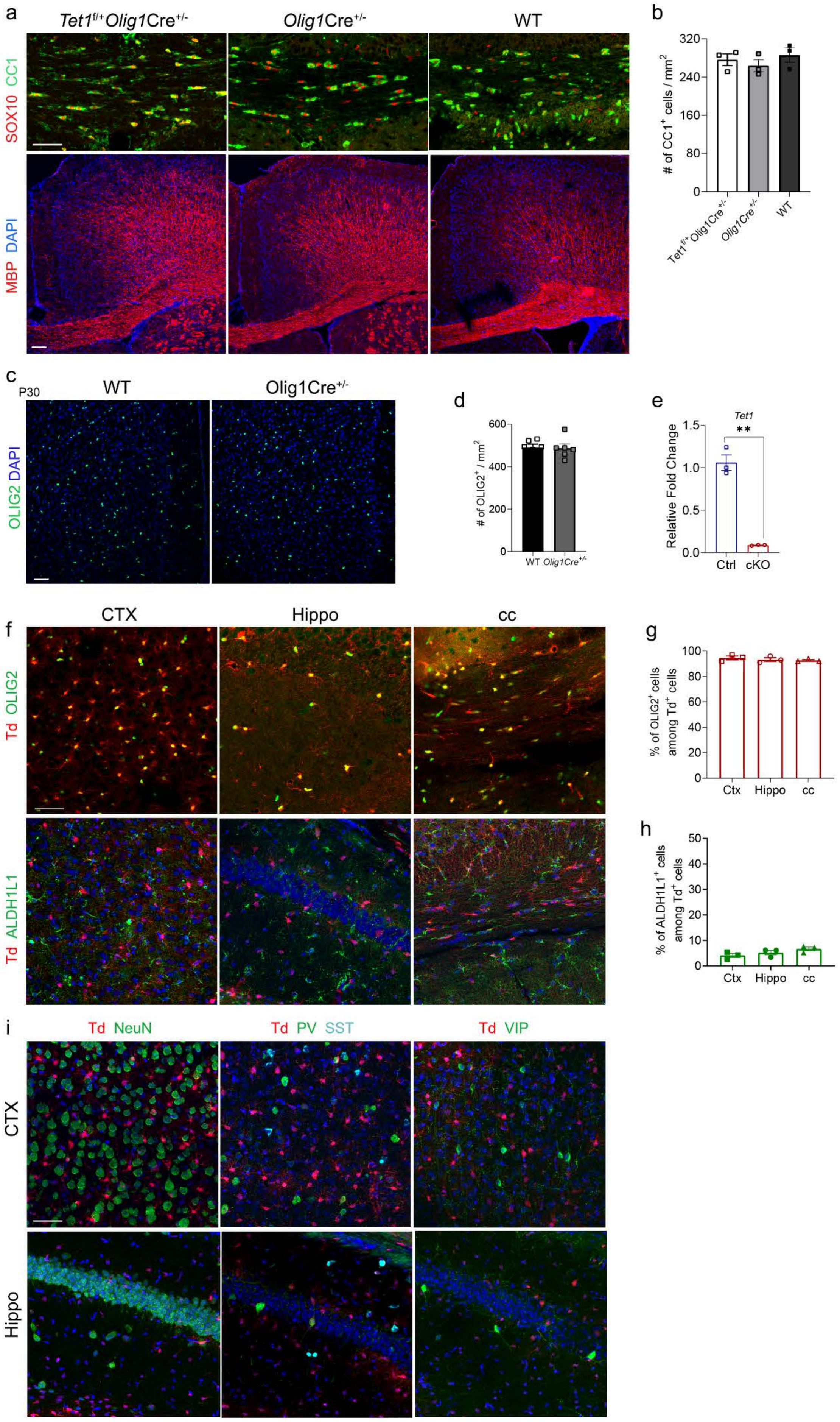
a
Tet1f/+Olig1Cre+/-
Olig1Cre+/-
WT
SOX10 CC1
MBP DAPI
b
### of CC1+ cells / mm2
320
240
160
80
0
Tet1f/+Olig1Cre+/-
Olig1Cre+/-
WT
c
P30
WT
Olig1Cre+/-
OLIG2 DAPI
d
### of OLIG2+ / mm2
600
400
200
0
WT
Olig1Cre+/-
e
Tet1
**
Relative Fold Change
1.5
1.0
0.5
0.0
Ctrl
cKO
f
CTX
Hippo
cc
Td OLIG2
Td ALDH1L1
g
% of OLIG2+ cells among Td+ cells
100
80
60
40
20
0
Ctx
Hippo
cc
h
% of ALDH1L1+ cells among Td+ cells
50
40
30
20
10
0
Ctx
Hippo
cc
i
Td NeuN
Td PV SST
Td VIP
CTX
Hippo

**Supplementary Fig. 3 OL differentiation and myelination are similar among *Tet1* heterozygous mice, Cre control, and wild-type mice, and *Olig1*-Cre is predominantly restricted to OL lineage**

(a) Representative immunostaining of P14 brains from indicated mice for mature OL marker CC1, OL lineage marker SOX10, and myelin protein MBP. Scale bar, 50 μm.

(b) Quantification of $CC1^+$ cells in P14 brains of indicated mice. Data are Means ± SEM ($n$ = 3 animals per group). One way ANOVA, $F_{(2,6)} = 0.688$, $p = 0.538$.

(c) Representative immunostaining of P30 cortex from indicated mice for OLIG2. Scale bar, 50 μm.

(d) Quantification the density of $OLIG2^+$ cells reveal no significant difference between indicated mice. Data are Means ± SEM ($n$ = 6 animals per group). Two-tailed unpaired $t$ test, $t = 0.076$, $df = 10$, $p = 0.941$.

(e) Real-time PCR quantification of *Tet1* mRNA in OPCs purified from *Tet1* cKO brain. Data are Means ± SEM ($n$ = 3 independent cultures each performed in triplicate). Two-tailed unpaired separate variance estimation $t$ test, **, $t = 10.61$, $df = 2.005$, $p = 0.0087$.

(f) Representative immunostaining of OLIG2 and ALDH1L1 in cortex (CTX), Hippocampus (Hippo) and corpus callosum (cc) in P21 *Olig1*Cre-tdTomato mice. Scale bar, 50 μm.

(g) Quantification the percentage of $OLIG2^+$ cells among $Td^+$ cells from different brain region. Data are Mean ± SEM ($n$ = 3 animals each group). One way ANOVA, $F_{(2,6)} = 4.465$, $p = 0.442$.

(h) Quantification the percentage of $ALDH1L1^+$ cells among $Td^+$ cells from different brain region. Data are Mean ± SEM (n=3 animals each group). Kruskal-Wallis H test, $\chi^2$=3.200, $df$ =2, $p$ = 0.202.

(i) Representative immunostaining of NeuN, PV, SST and VIP in P21 cortex and hippocampus of *Olig1*Cre-tdTomato mice. Rare co-labeling of NeuN, PV, SST and VIP with Td was observed. Scale bar, 50 μm.

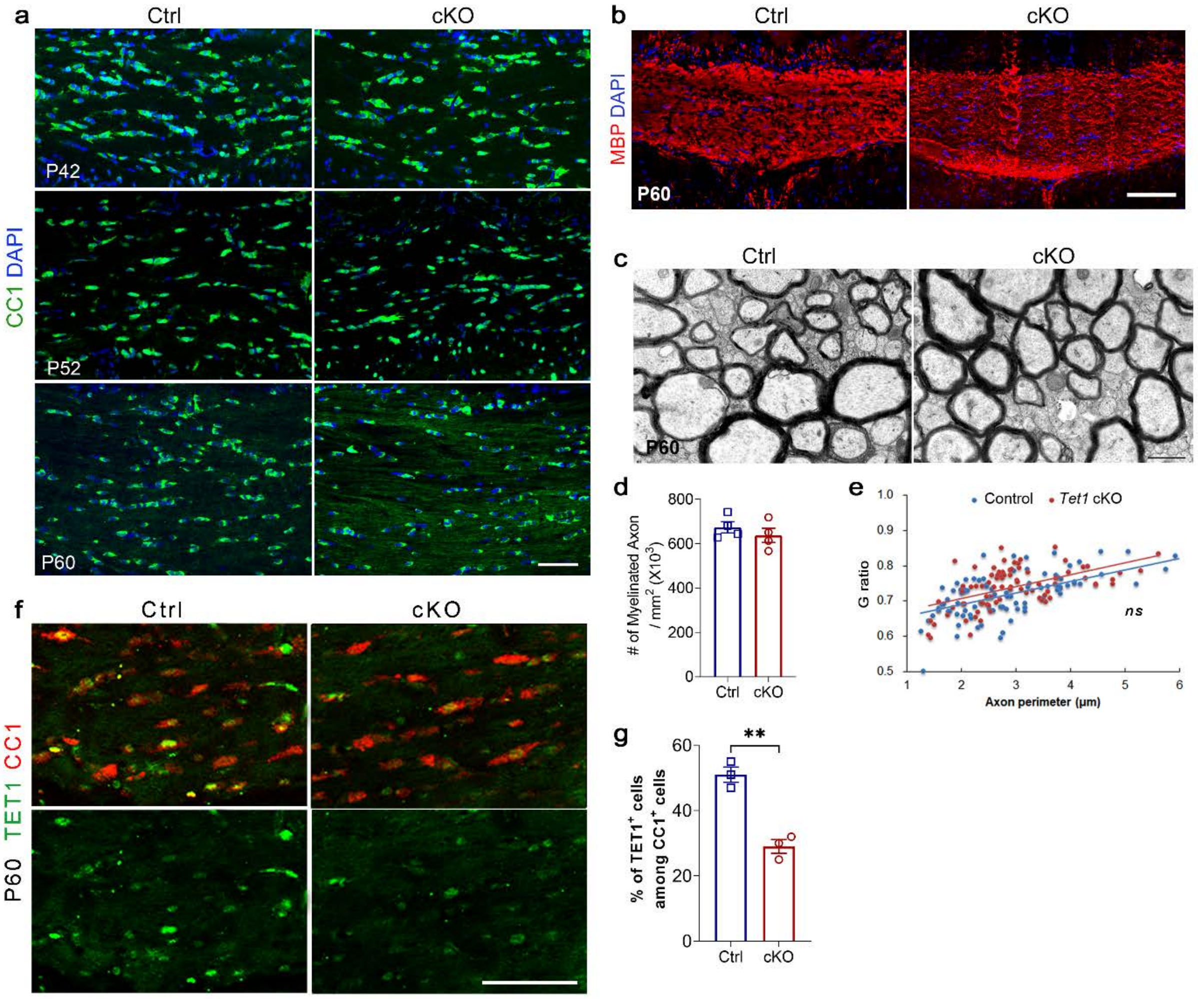


**Supplementary Fig. 4 Normal myelination in adult *Tet1* cKO mice**

(a) Representative immunostaining images of corpus collosum from P42, P52 and P60 controls and *Tet1* cKO mice stained for CC1. Scale bar, 50 μm.

(b) Representative immunostaining images of corpus collosum from P60 controls and *Tet1* cKO mice stained for MBP. Scale bar, 50 μm.

(c) Representative electron micrographs of corpus collosum from control and *Tet1* cKO mice at P60. Scar bar, 0.5 μm.

(d) Quantification the numbers of myelinated axons in defined areas from P60 control and *Tet1* cKO mice. Data are Means ± SEM ($n$ = 3 animals per group). Two-tailed unpaired $t$ test, $t$ = 0.0891, $df$ = 6, $p$ = 0.407.

(e) G ratios versus axonal perimeters for control and *Tet1* cKO mice at P60. *ns*; not significant, Friedman M test, $\chi^2$ = 2.390, $df$ = 1, $p_{\text{between group}}$ = 0.122 (> 80 myelinating axon counts from 3 animals each genotype).

(f) Representative images for TET1 and CC1 expression in corpus callosum of P60 mice. Scale bar, 50 μm.

(g) Quantification of $TET1^+$ cells among $CC1^+$ cells in control and *Tet1* cKO mice. Data are presented as Mean ± SEM ($n$ = 3 animals per group). **, Two-tailed unpaired $t$ test, $t$ = 7.706, $df$ = 6, $p$ = 0.002.

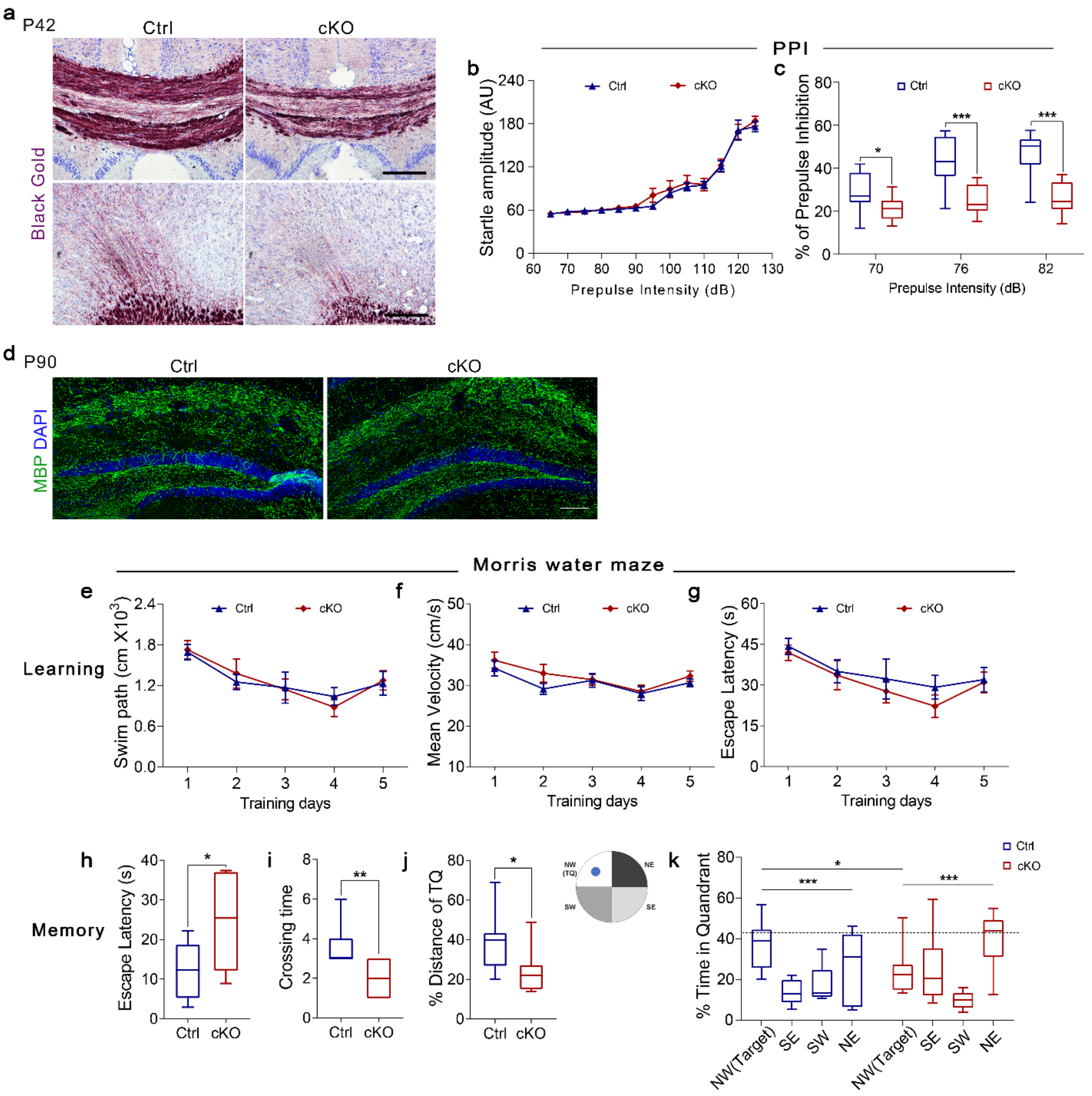


**Supplementary Fig. 5 Behavioral deficiency in *Tet1* cKO mice**

(a) Black gold staining shows myelin deficiency in P42 *Tet1* cKO mice. Scar bar, 100 μm.

(b) Similar baseline startle response was observed at multiple pulse intensities ranging from 65 to 125 dB in *Tet1* cKO mice and control littermates. Data were expressed as Mean ± SEM ($n$ = 13 for control mice and $n$ = 14 for *Tet1* cKO mice). RMANOVA, $F_{\text{between group }(1,25)} = 0.1836$, $p$ = 0.6720, $F_{\text{within group }(12,300)} = 144.6$, $p$ <0.0001. AU, arbitrary units.

(c) Impaired PPI in *Tet1* cKO mice for 70, 76 and 82 dB prepulse sound level. Boxes were expressed as quartiles (median, 25$^{th}$ and 75$^{th}$ percentile), whiskers were defined as minima and maxima ($n$ = 13 for control mice and $n$ =14 for *Tet1* cKO mice). Two-tailed unpaired $t$ test, 70dB: $t$ = 2.724, $df$ = 25, $p$ = 0.012; 76dB: $t$ = 4.913, $df$ = 25, $p$ = 0.000047; 82dB: $t$ = 5.611, $df$ = 25, $p$ = 0.000008.

(d) Representative MBP immunostaining for P90 adult mice in the hippocampus reveals comparable myelin between groups. Scar bar, 100 μm.

(e-g) Working memory impairment in *Tet1* cKO adult mice was revealed in the Morris water maze. *Tet1* mutant show normal spatial reference memory acquisition during 5-day training, indicated by the swimming distance (e), the mean swimming velocity (f) and the escape latency to the hidden platform (g). Data are expressed as Mean ± SEM ($n$ = 8 animals per group, four trials per day). RMANOVA test: (e) $F_{\text{between group }(1,14)} = 0.001$, $p$ = 0.976, $F_{\text{within group }(4,56)} = 7.260$, $p$ = 0.000088;

(f) $F_{\text{between group }(1,14)} = 1.100$, $p = 0.312$, $F_{\text{within group }(4,56)} = 8.034$, $p = 0.000034$; (g) $F_{\text{between group }(1,14)} = 0.765$, $p = 0.397$, $F_{\text{within group }(4,56)} = 6.571$, $p = 0.000208$.

(h-k) Probe trial for short-term memory retention was carried out 24hr after the last training. Compared to the control group, *Tet1* cKO mice spent longer time in searching the removed platform position (h) and crossed less times through the position (i). The reduced swimming distance (j) and swimming time in the target quadrants (TQ) indicate impaired spatial reference memory retention in *Tet1* mutant (k). NW, northwest; SE, southeast; SW, southwest; NE, northeast. Boxes were expressed as quartiles (median, $25^{th}$ and $75^{th}$ percentile), whiskers were defined as minima and maxima ($n = 8$ animals per group). (h): *, Two-tailed unpaired *t* test, $t = -2.650$, $df = 14$, $p = 0.019$; (i): **, Mann-Whitney U test, $z = -2.711$, $p = 0.007$; (j): *, Mann-Whitney U test, $z = -2.205$, $p = 0.027$; (k): Two-tailed unpaired *t* test for MW quadrant between Ctrl and cKO group, *, $t = 2.436$, $df = 14$, $p = 0.029$. One way ANOVA for % time in different quadrant. Ctrl: ***, $F_{(3,28)} = 8.153$, $p = 0.000467$. NW vs. SE, ***, $q = 24.4375$, $p = 0.000461$; NW vs. SW, **, $q = 19.58000$, $p = 0.005$; NW vs. NE, $q = 10.8500$, $p = 0.198$; SE vs. SW, $q = 4.85750$, $p = 0.798$; SE vs. NE, $q = -13.58150$, $p = 0.073$; SW vs. NE, $q = -8.73000$, $p = 0.372$. cKO: ***, $F_{(3,28)} = 7.685$, $p = 0.00067$. NW vs. SE, $q = -0.47375$, $p = 1.0$; NW vs. SW, $q = 14.54850$, $p = 0.116$; NW vs. NE, $q = -15.47500$, $p = 0.086$; SE vs. SW, $q = 15.01625$, $p = 1.0$; SE vs. NE, $q = -15.00125$, $p = 1.0$; SW vs. NE, ***, $q = -30.01750$, $p = 0.000266$.

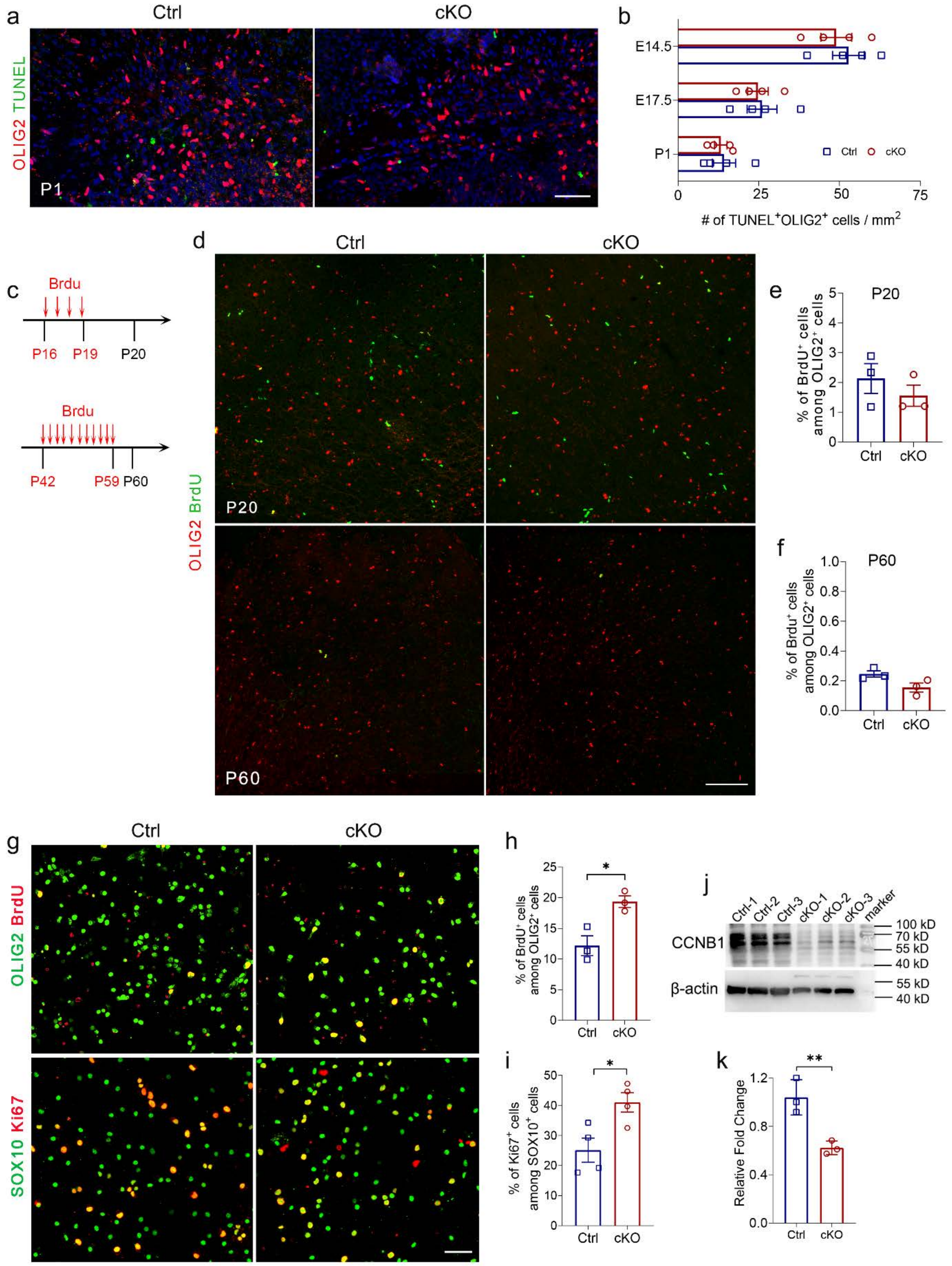


**Supplementary Fig. 6 Normal OPC apoptosis and increased proliferation in OPCs purified from *Tet1* cKO mice**

(a) Representative images of P1 control and *Tet1* cKO brains stained for OLIG2 after TUNEL staining. Scale bar, 50 μm.

(b) Quantification of TUNEL$^{+}$ OLIG2$^{+}$ cells in control and *Tet1* cKO brains from E14.5 to P1 stages. Data are Means ± SEM ($n$ = 4 animals each group). No significant differences were observed.

Two-tailed unpaired *t* test, E14.5: $t = 0.246$, $df = 6$, $p = 0.714$; E17.5: $t = 0.223$, $df = 6$, $p = 0.831$; P1: $t = 0.542$, $df = 6$, $p = 0.604$.
(c) Diagram showing BrdU injection to label proliferating cells in P20 and P60 mice.
(d) Representative images of BrdU and OLIG2 immunostaining in P20 and P60 cortex. Scale bar, 100 μm.
(e-f) Quantification the percentage of $BrdU^+$ cells among $OLIG2^+$ cells, suggesting no difference between control and *Tet1* cKO mice at indicated stages. Data are Means ± SEM ($n = 3$ animals each group). (e): Mann-Whitney U test, $z = -0.655$, $p = 0.513$; (f): Two-tailed unpaired *t* test, $t = 2.444$, $df = 4$, $p = 0.071$.
(g) Representative images of double immunostaining OPC cultures with BrdU / OLIG2, or Ki67 / SOX10 to illustrate the proliferating cells in OL lineage. Scale bar, 50 μm.
(h-i) Comparison the proportion of $BrdU^+$ (h) and $Ki67^+$ cells (i) among OPCs between control and *Tet1* cKO mice. Note the increased number of proliferating OPCs in *Tet1* cKO group. Data are Mean ± SEM ($n = 3$ independent experiments each group). (h): *, Two-tailed unpaired *t* test, $t = -3.832$, $df = 4$, $p = 0.019$; (i): *, Two-tailed unpaired *t* test, $t = -3.093$, $df = 6$, $p = 0.021$.
(j) Western blot assay for the expression of CCNB1 in purified control and *Tet1* cKO OPC cultures. β-actin is used for loading control.
(k) Histogram shows fold changes measured by densitometry in *Tet1* cKO group relative to control after normalization to β-actin levels. Data are Mean ± SEM ($n = 3$ independent cultures). **, Two-tailed unpaired *t* test, $t = 4.638$, $df = 4$, $p = 0.0098$.

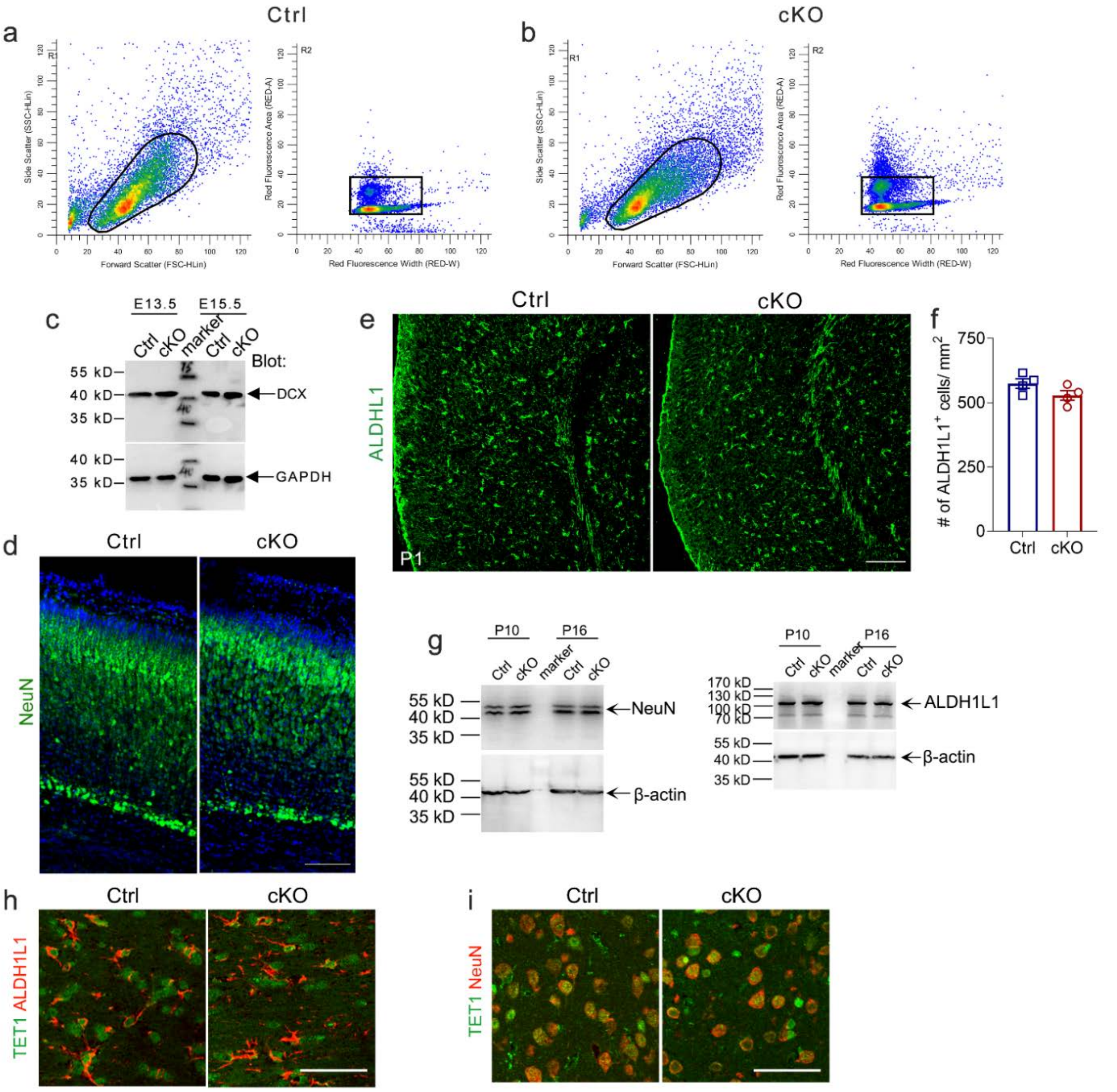


**Supplementary Fig. 7 Normal astrocytes and neurons development in *Tet1* cKO mice**

(a-b) Representative gating strategies for flow-cytometry analysis of cell cycle in purified OPCs from Control (a) and *Tet1* cKO (b) mice. The sorting strategy is as follows: on the first sort, the major cell groups is included by drawing a gate around cells with 1.0-7.5 K forward (FSC-HLin) and side (SSC-HLin) scatter. On the second sort, the single cell was gated in 3.0K-5.0K Red Fluorescence Width and 1.5K-6.0K Area. By looking at forward scatter vs width, we exclude cellular debris and large cell masses.

(c) Western blot assay for expression of DCX in control and *Tet1* cKO E13.5 and E15.5 brains. GAPDH was detected as a loading control.

(d) Representative images of P1 control and *Tet1* cKO brains immunostained for NeuN. Scale bar, 100 μm.

(e) Representative images of P1 control and *Tet1* cKO S1 cortex stained for ALDH1L1. Scale bar, 100 μm.

(f) Quantification of ALDH1L1$^{+}$ cells from P1 control and *Tet1* cKO S1 cortex. Data are Means ± SEM ($n$ = 3 animals per group). Two-tailed unpaired $t$ test, $t$ = 1.755, $df$ = 6, $p$ = 0.130.

(g) Western blot assay for the expression of ALDH1L1 and NeuN in the brain samples from postnatal control and *Tet1* cKO mice; β-actin was detected as a loading control.

(h-i) Representative images of P10 control and *Tet1* cKO brains immunostained for TET1/ALDH1L1 (h) and TET1/NeuN (i). Scale bar, 50 μm.

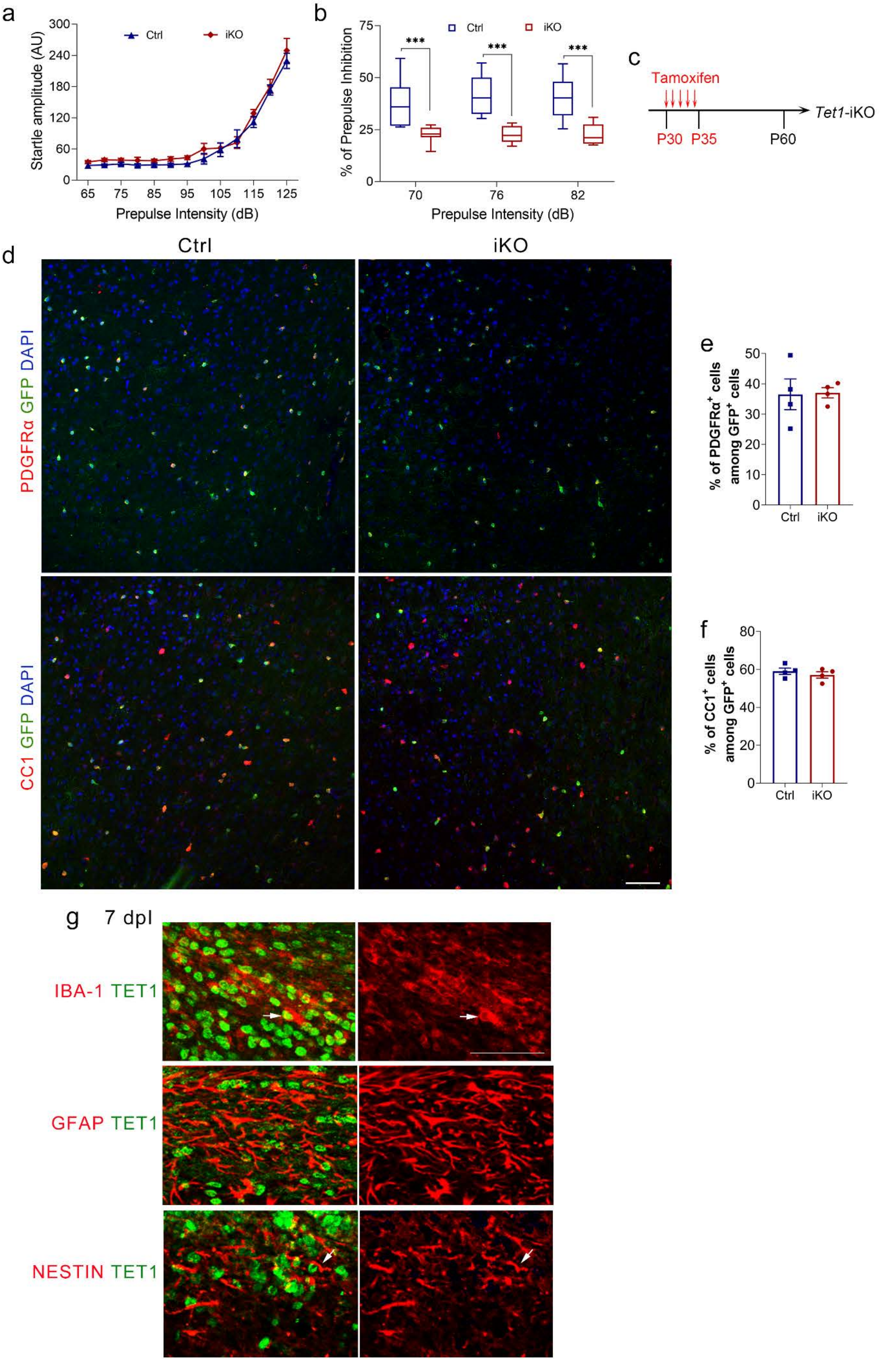
a
Ctrl
iKO
Startle amplitude (AU)
Prepulse Intensity (dB)
b
% of Prepulse Inhibition
***
70
76
82
Prepulse Intensity (dB)
c
Tamoxifen
P30
P35
P60
Tet1-iKO
d
Ctrl
iKO
PDGFRα GFP DAPI
CC1 GFP DAPI
e
% of PDGFRα+ cells among GFP+ cells
f
% of CC1+ cells among GFP+ cells
g
7 dpl
IBA-1 TET1
GFAP TET1
NESTIN TET1

**Supplementary Fig. 8 Impaired PPI in *Tet1* iKO (*NG2-CreER; Tet1$^{flox/flox}$*) mice and OPC fate in adult *Tet1* iKO-GFP mice**

(a) Similar baseline startle response was observed at multiple pulse intensities ranging from 65 to 125 dB in *Tet1* iKO mice and control littermates. Data were expressed as Mean ± SEM ($n$ = 8 for *Tet1* cKO and control mice). RMANOVA, $F_{\text{between group }(1,14)} = 3.589$, $p = 0.079$, $F_{\text{within group }(2.747,41.21)} = 458.2$, $p < 0.0001$. AU, arbitrary units.

(b) Impaired PPI in *Tet1* iKO mice for 70, 76 and 82 dB prepulse sound level. Boxes were expressed as quartiles (median, 25$^{th}$ and 75$^{th}$ percentile), whiskers were defined as minima and maxima ($n$ = 11 for *Tet1* iKO and control mice). ***, Two-tailed unpaired separate variance estimation $t$ test, 70dB: $t = 4.584$, $df = 12.804$, $p = 0.00053$; 76dB: $t = 6.447$, $df = 13.771$, $p = 0.000017$; 82dB: $t = 5.386$, $df = 14.432$, $p = 0.000086$.

(c) Diagram showing Tamoxifen administration to induce Cre recombination from P30 in *Tet1* iKO mice with GFP Cre reporter. Mice were sacrificed for analysis at P60.

(d) Representative double immunostaining for PDGFRα / GFP and CC1 / GFP in the cortex of P60 *Tet1* iKO, R26-YFP mice. Scale bar, 100 μm.

(e) Quantification the percentage of PDGFRα$^{+}$ cells in GFP$^{+}$ cells in control and *Tet1* iKO mice. Data are Mean ± SEM ($n$ = 4 of animals each group). Two-tailed unpaired $t$ test, $t = -0.102$, $df = 6$, $p = 0.922$.

(f) Quantification the percentage of CC1$^{+}$ cells in GFP$^{+}$ cells in control and *Tet1* iKO mice. Data are Mean ± SEM ($n$ = 4 of animals each group). Two-tailed unpaired $t$ test, $t = 0.827$, $df = 6$, $p = 0.440$.

(g) TET1 expression can be detected in microglia and NPCs, not in astrocytes, in LPC lesion site. Representative double immunostaining of TET1 with microglia marker IBA1, astrocyte marker GFAP and neural progenitor maker NESTIN in the lesion site. TET1 expression in IBA-1$^{+}$ cells and few NESTIN$^{+}$ cells are indicated by the arrows. Scale bar, 100 μm.

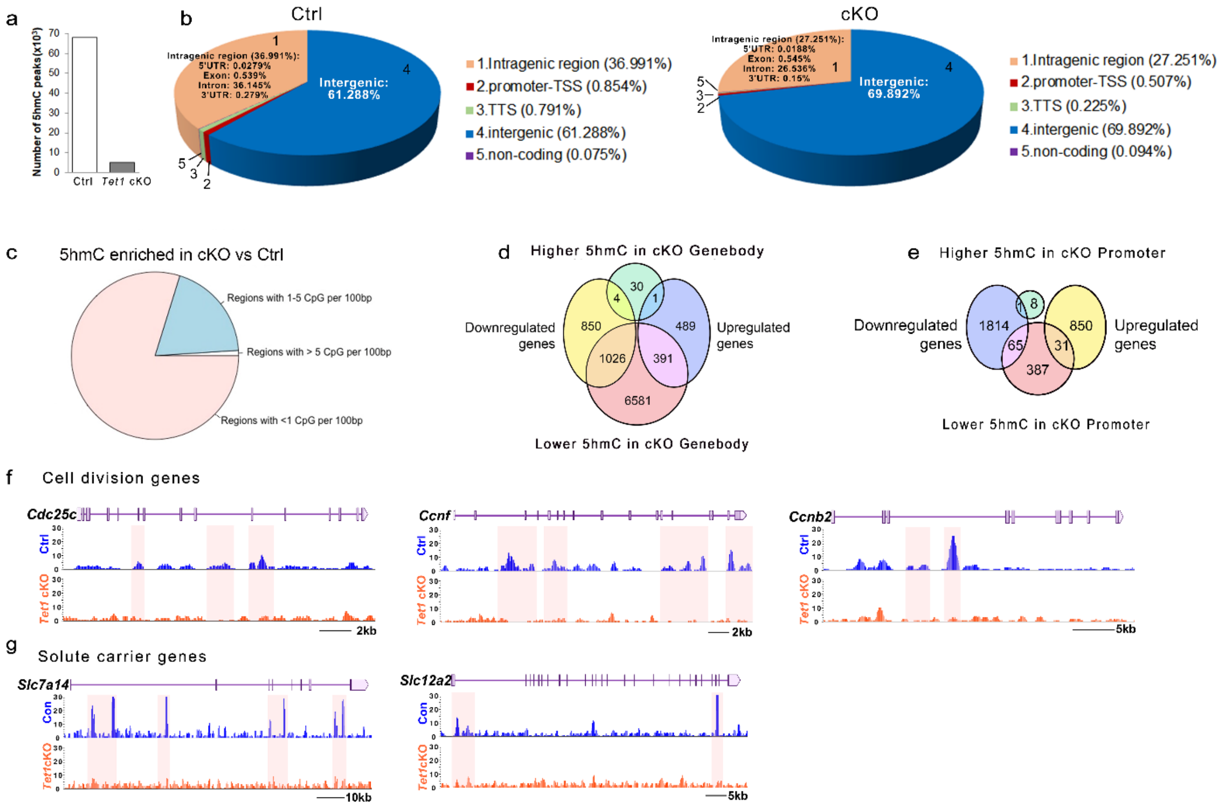


**Supplementary Fig. 9 5hmC distribution in the genome of OPCs is altered in *Tet1* cKO mice**

(a) The number of total 5hmC peaks from hMeDIP-seq in control and *Tet1* cKO mice.
(b) Genomic distribution of 5hmC in OPCs from control and *Tet1* cKO mice.
(c) Proportion of differentially hydroxymethylated regions with different CpG densities.
(d) Venn diagram of the overlap between differentially hydroxymethylated peaks at gene bodies and differentially expressed genes.
(e) Venn diagram of the overlap between genes differentially hydroxymethylated at promoters and differentially expressed genes.
(f) Representative 5hmC peaks of cell division genes in OPCs derived from control (blue) and *Tet1* cKO (orange) mice.
(g) Representative 5hmC peaks of *Slc* family genes in OPCs derived from control (blue) and *Tet1* cKO (orange) mice.

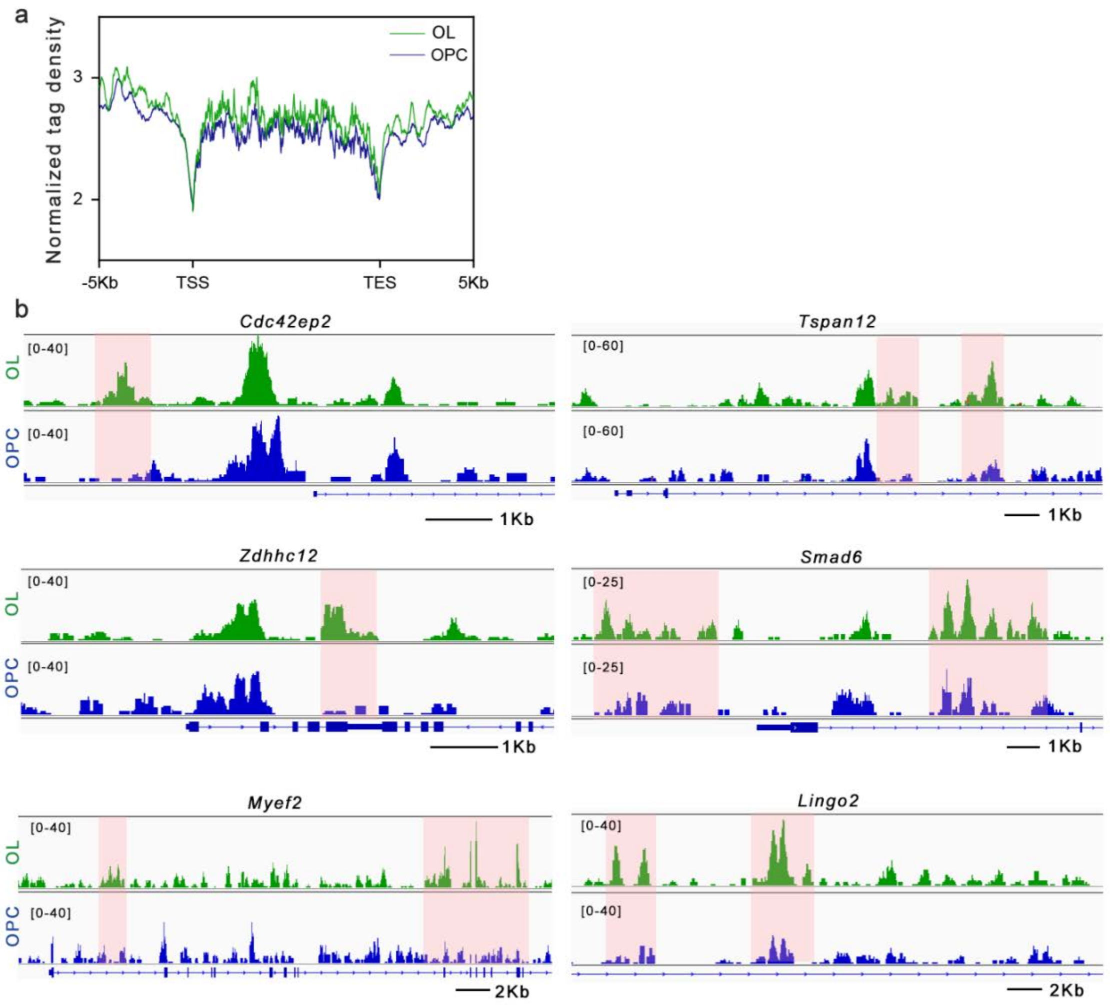


**Supplementary Fig. 10 Comparation 5hmC land scape between OPC and OL.**
(a) Normalized 5hmC tag density distribution from normal OPCs and OLs.
(b) Snapshots of 5hmC profiles of representative genes with hyper-hydroxyemethylated regions in OLs (green) compared with OPCs (blue). Pink boxes label the hyper-5hmC region in OLs.

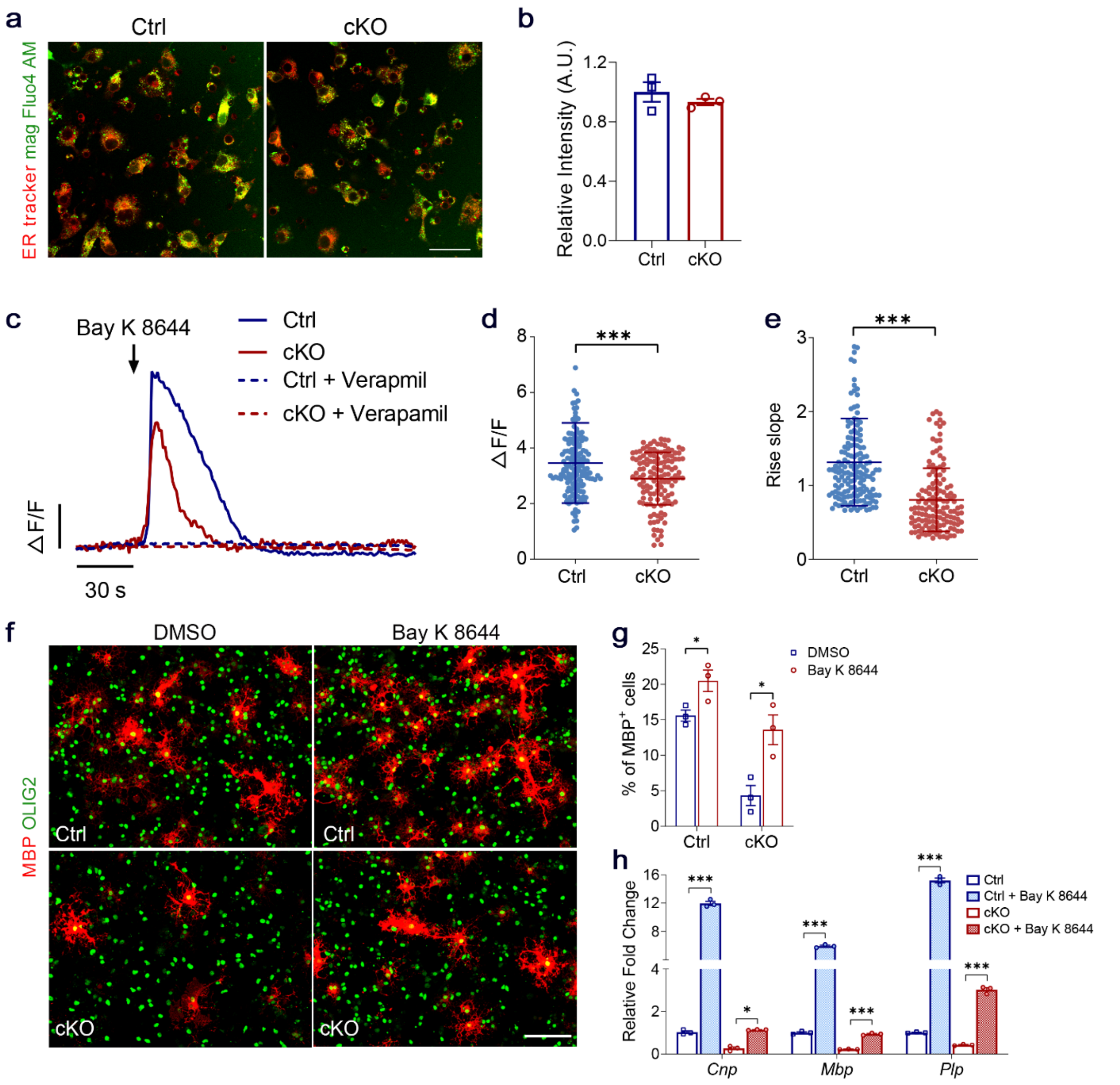


**Supplementary Fig. 11 Activating calcium rescues the impaired differentiation in *Tet1* depleted OPCs**

(a) Representative images of living OPCs from control and *Tet1* cKO mice loaded with mag Fluo4 AM (Green) and ER tracker (Red). Scale bar, 40 μm.

(b) Relative fluorescence intensity of mag Fluo4 AM measured by spectrofluorometer from control and *Tet1* cKO OPC cultures. Data are Means ± SEM ($n$ = 3 independent cultures), Two-tailed unpaired $t$ test, $t$ = 1.933, $df$ = 4, $p$ = 0.125.

(c) Representative traces of Fluo4 intensity of OPCs from control and *Tet1* cKO mice following application of Bay K 8644 and Verapamil.

(d) Amplitude changes ($\Delta F / F$) after 10 μM Bay K 8644 treatment of control and *Tet1* cKO OPCs. Data are Means ± SD ($n$ = 157 cells from three independent cultures in Ctrl group, $n$ = 130 cells from three independent cultures in cKO group). ***, Two-tailed unpaired $t$ test, $t$ = 3.847, $df$ = 285, $p$ = 0.0001.

(e) The average rise in slope of Fluo4 intensity after Bay K 8644 addition in control and *Tet1* cKO OPCs. Data are Means ± SD ($n$ = 157 cells from three independent cultures in Ctrl group, $n$ = 130 cells from three independent cultures in cKO group). ***, Two-tailed unpaired separate variance estimation $t$ test, $t$ = 8.409, $df$ = 280.3, $p$ < 0.0001.

(f) Representative images of MBP / OLIG2 immunostaining of Bay K 8644-treated control and *Tet1* cKO OPCs. Scar bar, 100 μm.

(g) Percentage of $MBP^+$ cells among $OLIG2^+$ cells after 3 days of Bay K 8644 treatment in control

and *Tet1* cKO OPCs. Data are Means ± SEM ($n$ = 3 of independent cultures each group). Two-tailed unpaired $t$ test, Ctrl: *, $t$ = -2.877, $df$ = 4, $p$ = 0.045; cKO: *, $t$ = -3.655, $df$ = 4, $p$ = 0.022.
(h) Quantitative real-time PCR of *Cnp*, *Mbp*, and *Plp* after three days of Bay K 8644 treatment in control and *Tet1* cKO OPCs relative to vehicle treatment. Data are Means ± SEM of transcript levels after normalization from $n$=3 independent experiments each performed in triplicate. *Cnp*: One way ANOVA, $F_{(3,8)}$ = 1278, $p$ < 0.0001; Ctrl vs. Ctrl+Bay K 8644: ***, $q$ = 69.94, $df$ = 8, $p$ < 0.0001; cKO vs. cKO+Bay K 8644: *, $q$ =5.594, $df$ = 8, $p$ = 0.0177. *Mbp*: One way ANOVA, $F_{(3,8)}$ = 1436, $p$ < 0.0001; Ctrl vs. Ctrl+Bay K 8644: ***, $q$ = 71.04, $df$ = 8, $p$ < 0.0001; cKO vs. cKO+Bay K 8644: ***, $q$ =10.34, $df$ =8, $p$ = 0.0004. *Plp*: One way ANOVA, $F_{(3,8)}$ = 1530, $p$ <0.0001; Ctrl vs. Ctrl+Bay K 8644: ***, $q$ = 79.90, $df$ = 8, $p$ <0.0001; cKO vs. cKO+Bay K 8644: ***, $q$ =14.61, $df$ =8, $p$ <0.0001.

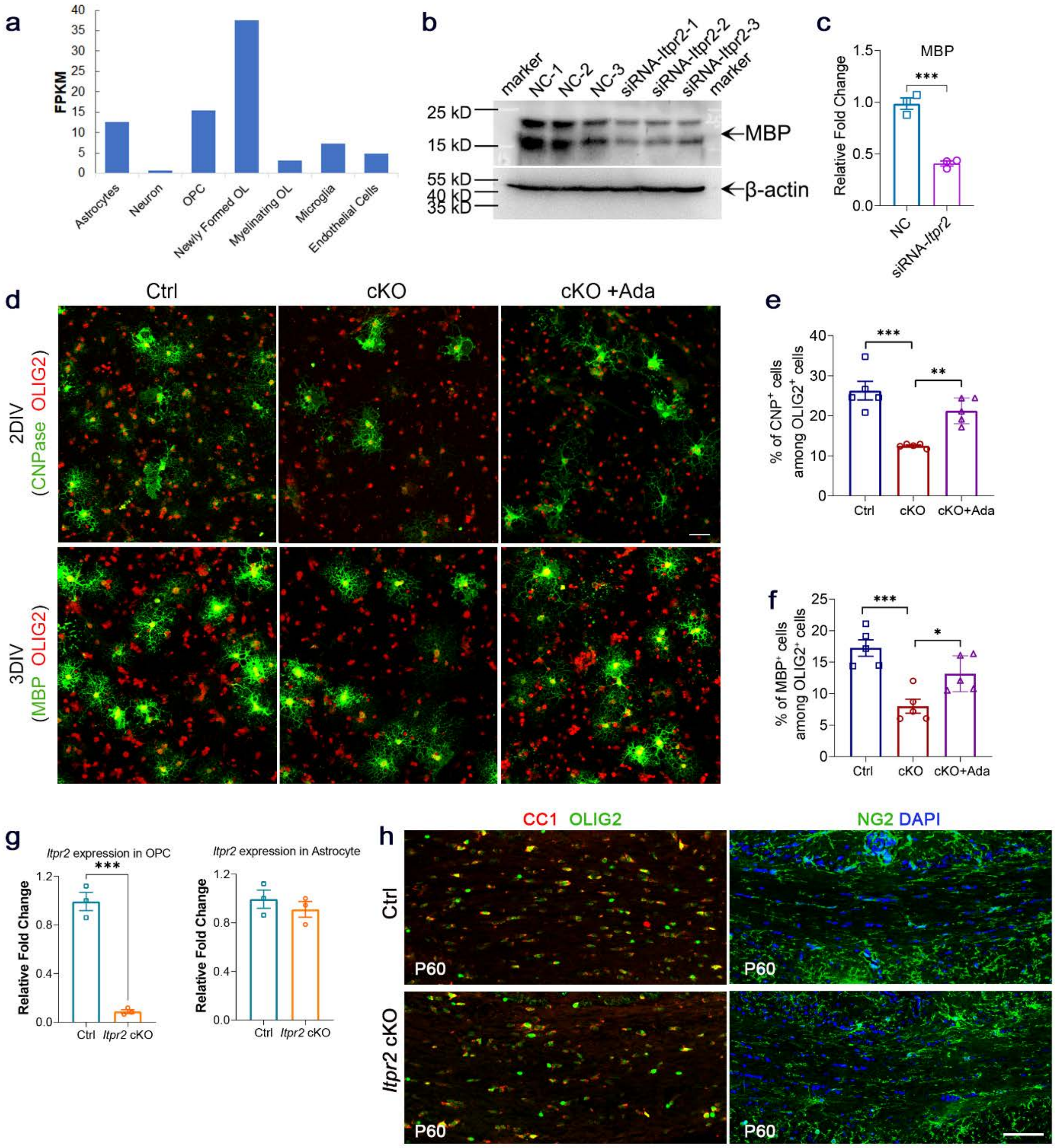


**Supplementary Fig. 12 ITPR2, one of TET1-5hmC target genes, is required for proper OPC differentiation**

(a) Column diagram shows the expression of ITPR2 in different neural cell types. The diagram is made from data in http://jiaqianwulab.org/resource.htm.

(b) Western blot assay for MBP showed decreased expression in siRNA-*Itpr2* transfected OPC cultures after three days in differentiation medium.

(c) Histogram shows MBP expression fold change measured by densitometry in siRNA-*Itpr2* group relative to control after normalization to β-actin levels. Data are Means ± SEM ($n$ = 3 transfections). ***, Two-tailed unpaired $t$ test, $t$ = 9.388, $df$ = 4, $p$ = 0.000717.

(d) Representative images of CNPase, MBP and OLIG2 immunostaining in control, *Tet1* cKO and Ada-treated *Tet1* cKO OPCs at 2DIV and 3DIV. Scale bar, 50 μm.

(e) Percentage of $CNP^+$ cells among $OLIG2^+$ cells after 2 days of Ada treatment in *Tet1* cKO OPCs. Data are Means ± SEM ($n$ = 4 of independent cultures each group). One way ANOVA, $F_{(2,12)}$ = 19.422, $p$ = 0.000173. Tukey's multiple comparisons test, Ctrl vs. cKO: ***, $q$ = 8.708, $df$ =

12, $p$ = 0.0001, Ctrl vs. cKO+Ada: $q$ = 3.169, $df$ = 12, $p$ = 0.1042, cKO vs. cKO+Ada: **, $q$ = 5.539, $df$ = 12, $p$ = 0.0054.

(f) Percentage of $MBP^+$ cells among $OLIG2^+$ cells after 3 days of Ada treatment in *Tet1* cKO OPCs. Data are Means ± SEM ($n$ = 5 of independent cultures each group). One way ANOVA, $F_{(2,12)}$ = 14.058, $p$ = 0.000717. Tukey's multiple comparisons test, Ctrl vs. cKO: ***, $q$ = 7.485, $df$ = 12, $p$ = 0.0005; Ctrl vs. cKO+Ada: $q$ = 3.306, $df$ = 12, $p$ = 0.0886, cKO vs. cKO+Ada: *, $q$ = 4.180, $df$ = 12, $p$ = 0.0300.

(g) qRT-PCR assay tests the mRNA expression of *Itpr2* in OPCs and astrocytes purified from control and *Itpr2* cKO mice. Data are presented as Mean ± SEM of transcript levels after normalization from $n$=3 independent experiments each performed in triplicate. Two-tailed unpaired $t$ test, OPC: ***, $t$ = 11.794, $df$ = 4, $p$ = 0.000296; Astrocyte: $t$ = 0.852, $df$ = 4, $p$ = 0.442.

(h) Representative immunostaining of CC1 / OLIG2 and NG2 in corpus callosum of P60 mice. Scale bar, 50 μm.

**Supplementary Table1. Summary of 5hmC peak region, primer information and results for BisulPlus™ Loci 5hmC-qPCR assay in certain genes.**

| | Gene Name (Chr #) | 5hmC peak region | Hydroxymethylated primers | Un-hydroxymethylated primers | Relative 5hmC abundance | p value |
|---|---|---|---|---|---|---|
| OPC associated genes | *Cspg4* (Chr9) | 56861264-56862568 | TTTAGATAGAAAAGTATTTGTCGGG<br>TTTAGATAGAAAAGTATTTGTCGGG-3 | TTAGATAGAAAAGTATTTGTTGGGG<br>AATTTCCCTTAACTTAAATCAACCA | 2.576 | 0.0148 |
| | *Slc22a3* (Chr17) | 12498956-12503627 | ATTATATCGACGAAGAGGAAGGC<br>CAAACACTAAAAAAAACGAACGAAT | TCCAGACTCTCCCTGCTTAAC<br>TCTTCTCTCCTTCTGCCACTG | 41.810 | <0.0001 |
| | *Wnt10a* （Chr1） | 74792174-74793286 | TCCAGACTCTCCCTGCTTAAC<br>TCTTCTCTCCTTCTGCCACTG | TTATATTGATGAAGAGGAAGGTGAAG<br>CCAAACACTAAAAAAAACAAACAAAT | 2.011 | <0.0001 |
| iOL associated genes | *Kndc1* (Chr7) | 139893099-139894079 | TAGGTTGAGATGTATTAGTTGTGGC<br>AATACTTTCCTCTAATTAACGCTCA | GAGATGTATTAGTTGTGGTGTGT<br>AATACTTTCCTCTAATTAACACTCA | 17.010 | 0.0322 |
| | *Olfr279* （Chr15） | 98496136-98496853 | TTAGGAAAATAAATAATTTTCGAGA<br>AAAAACCAAAACATCTACCCTACG | TTAGGAAAATAAATAATTTTTGAGA<br>AAAAAACCAAAACATCTACCCTACA | 1.686 | 0.0187 |
| | *Zdhhc12* （Chr2） | 30093033-30093493 | GAGTAGGGTAATTATTAACGCGAAG<br>ACCCTAAAAACTATAACTCCGATCC | GGAGTAGGGTAATTATTAATGTGAAG<br>TCCAACCCTAAAAACTATAACTCCA | 1.179 | 0.0412 |
| mOL associated genes | *Mag* (Chr7) | 30900452-30901587 | TAATGGTAATTAGGATGGTAAAGGC<br>CCCTAAAAACTCCTAATCTCGTTCT | TGGTAATTAGGATGGTAAAGGTGAT<br>CCCTAAAAACTCCTAATCTCATTCT | 3.170 | <0.0001 |
| | *Elovl7* (Chr13) | 108215666-108216506 | GGTTTCGAATTTTTTGTGTAGTTTC<br>CTAACAAACATAAAACCTCCGATTT | TTTGAATTTTTTGTGTAGTTTTGGAT<br>CTAACAAACATAAAACCTCCAATTT | 9.007 | 0.0383 |
| | *Arl2* （Chr19） | 6139651-6140428 | TATTTTGTTAGGTAGGTTTTCGAAG<br>ACTAAAACCCAAATATAAACGCTTA | TATTTTGTTAGGTAGGTTTTTGAAG<br>ACTAAAACCCAAATATAAACACTTA | 1.641 | 0.0346 |

| | | | | | | |
|---|---|---|---|---|---|---|
| | *Cdc42ep2*<br>（Chr19） | 5924966-<br>5926196 | AATCGAGGGAAATTTTTTATTGC<br>CAACCTATACCTAAAACCTCGAATT | AAATTGAGGGAAATTTTTTATTGTGT<br>CAACCTATACCTAAAACCTCAAATT | 3.086 | 0.0078 |
| Negative regulator | *Zfp28*<br>（Chr19） | 6387100-<br>6387562 | AATCGAGGGAAATTTTTTATTGC<br>CAACCTATACCTAAAACCTCGAATT | AAATTGAGGGAAATTTTTTATTGTGT<br>CAACCTATACCTAAAACCTCAAATT | 0.8052 | <0.0001 |
| | *Ngf*<br>(Chr3) | 102471240-<br>102471702 | AGTTTAGAGGGTAGTGTAGTCGTTT<br>GAAAACCGAATATAACCCGTAAC | AGTTTAGAGGGTAGTGTAGTTGTTT<br>ACAAACAAAAACCAAATATAACCCA | 0.1127 | 0.0489 |

**Supplementary Table 2. Transcriptome changes in OPCs from *Tet1* cKO mice**

| Upregualted gene | Log2 (cKO/OPC) | Downregulated gene | Log2 (cKO/OPC) |
|---|---|---|---|
| Ifi202b | 9.145461947 | Clca1 | -7.175680633 |
| Tlr1 | 7.216005351 | Ttr | -6.751171351 |
| Ccl6 | 6.581469776 | Cdh5 | -6.418979102 |
| Chrna10 | 5.859272075 | Gldn | -5.926284196 |
| Nts | 5.680285631 | Slc14a1 | -5.506635508 |
| Rprml | 5.460092219 | Serpina3n | -5.363509847 |
| Egr4 | 5.311048922 | Cyp3a13 | -5.303327196 |
| 4930539E08Rik | 5.191022921 | Klk6 | -5.200460001 |
| Sp140 | 5.138193072 | Gdnf | -5.165446393 |
| Cd74 | 5.115591761 | Cd34 | -4.980227037 |
| Clec4n | 5.01833284 | Mmp13 | -4.871453224 |
| Pf4 | 4.851203631 | Gprc5c | -4.854812179 |
| Olfr287 | 4.807492251 | Fam19a1 | -4.711317976 |
| Ankrd63 | 4.691102151 | Tmem196 | -4.710234354 |
| Gpr88 | 4.565407171 | Eps8l1 | -4.644114901 |
| Rpe65 | 4.50694568 | Ppl | -4.626566012 |
| Gabrd | 4.48625855 | Ptgfr | -4.600421631 |
| Ipcef1 | 4.302236827 | Alox5 | -4.592833791 |
| Fcgr2b | 4.301501803 | Slc5a7 | -4.576190491 |
| Rasgrp1 | 4.280273614 | Nek5 | -4.541786201 |
| Ccl7 | 4.232637713 | Dmp1 | -4.482922315 |
| Drd2 | 4.19944924 | Prl2c2 | -4.471212499 |
| Hist1h2ao | 4.129084596 | Tnn | -4.470936612 |
| Penk | 4.113720446 | Epcam | -4.384141741 |
| Cxcl3 | 4.083162402 | Akap14 | -4.376001719 |
| Acpp | 4.067836595 | Mobp | -4.367261996 |
| Gjb6 | 4.038321878 | Gm6588 | -4.338490814 |
| Tal1 | 3.967374033 | Tmem132c | -4.278063047 |
| Rxrg | 3.90493484 | Mag | -4.229839775 |
| Ddn | 3.877639332 | Akr1b8 | -4.22760776 |
| Il2rg | 3.828184135 | Gjc2 | -4.213118585 |
| Aldh1a1 | 3.77992922 | Thsd4 | -4.206536565 |
| Alox5ap | 3.771105764 | Cldn11 | -4.201819655 |
| Ucp2 | 3.719785286 | Lingo2 | -4.167865308 |
| Mx1 | 3.682453465 | Slc24a2 | -4.122341697 |
| Prdm8 | 3.682057339 | Syne3 | -4.054357343 |
| Wfdc17 | 3.668146027 | Tcea3 | -4.038440154 |
| Fpr2 | 3.647745314 | Tmprss5 | -4.033205429 |
| Matn4 | 3.639746697 | Gpr149 | -3.987163254 |
| Sstr3 | 3.625836727 | Slc9a2 | -3.982179637 |
| Vax2 | 3.624722319 | Foxd3 | -3.963006175 |
| Cd33 | 3.620399911 | Nipal1 | -3.951699209 |
| Saa3 | 3.614482374 | Tktl1 | -3.945976033 |
| Dclk3 | 3.614328493 | Speer4b | -3.929919951 |
| Hist1h2an | 3.566203981 | Lef1 | -3.925884721 |
| Dhrs7c | 3.54404446 | Ndst4 | -3.913697554 |
| Kcnh3 | 3.533685654 | Efhb | -3.907454539 |
| Ccl3 | 3.524430921 | Prima1 | -3.895949966 |
| Hapln1 | 3.524312164 | A930009A15Rik | -3.876244044 |
| Tmem210 | 3.517017083 | Mbp | -3.83788806 |
| Klk13 | 3.499657105 | Lpar1 | -3.807297249 |
| Pou6f2 | 3.499652366 | Arhgap28 | -3.804787829 |
| Nrsn2 | 3.465196646 | Il22ra1 | -3.799150889 |
| Camk2a | 3.447694419 | Sytl2 | -3.793151505 |
| Helt | 3.422689413 | Ano1 | -3.791471429 |

| | | | |
|---|---|---|---|
| Cacng3 | 3.416599066 | Ifi205 | -3.789019082 |
| Odf3l1 | 3.397542268 | Spag16 | -3.783930517 |
| Pla2g7 | 3.392131877 | Enpp6 | -3.775748763 |
| Slc6a7 | 3.377622996 | Tmprss3 | -3.775405949 |
| Ebf2 | 3.36749516 | Birc7 | -3.758199949 |
| Mal2 | 3.367493463 | Rec8 | -3.757412912 |
| Il1f9 | 3.357380671 | Adh7 | -3.753535519 |
| Camkv | 3.320219134 | Pappa2 | -3.752998563 |
| Susd3 | 3.298071882 | Gprc5a | -3.715098176 |
| Npr3 | 3.296655547 | Ppp1r36 | -3.688030186 |
| Nrgn | 3.279747676 | Gpr55 | -3.672814244 |
| Lcn2 | 3.26996332 | Tmem184a | -3.671877323 |
| Serpinb6b | 3.267353925 | Egfl6 | -3.667046112 |
| Hal | 3.262207906 | Nr1h4 | -3.654957232 |
| Slc39a12 | 3.25498555 | Nostrin | -3.654947293 |
| Ccr7 | 3.250056197 | Fndc3c1 | -3.654740402 |
| Rab37 | 3.225434877 | Tex22 | -3.647553772 |
| Gjd3 | 3.212815059 | Tmem125 | -3.632111551 |
| Rgs14 | 3.197068637 | Zfp92 | -3.609079751 |
| Lao1 | 3.194894621 | Plp1 | -3.606409287 |
| Hpcal4 | 3.184335316 | Kcne4 | -3.58134193 |
| Gm5796 | 3.181971517 | Sprr1a | -3.555737305 |
| Pou3f1 | 3.179737714 | Fam110c | -3.550938659 |
| Grem2 | 3.17488368 | Tmem163 | -3.54313798 |
| Htr1a | 3.173771425 | Ccl28 | -3.528091568 |
| Atp6v0d2 | 3.172856559 | Fhad1 | -3.501407616 |
| Asb16 | 3.172853144 | Trabd2b | -3.484577166 |
| Clic5 | 3.148546678 | Ccdc162 | -3.479230728 |
| Ppp1r3g | 3.141959174 | Trf | -3.4720488 |
| Hmgcs2 | 3.134186244 | Ccdc60 | -3.471533956 |
| Sowaha | 3.122734184 | Slc15a1 | -3.468361216 |
| Slc25a34 | 3.120880394 | Col8a1 | -3.457725791 |
| Nxnl2 | 3.097680554 | Igfals | -3.442750176 |
| Cd300ld | 3.095274999 | Akr1c21 | -3.429767736 |
| Slc35f4 | 3.088044553 | Prl2c3 | -3.424838901 |
| F13a1 | 3.08161526 | Tfpi2 | -3.423232885 |
| Tpo | 3.069685574 | Kndc1 | -3.421942274 |
| Gngt2 | 3.067836848 | Spp1 | -3.402946935 |
| C130060K24Rik | 3.045003338 | Snai2 | -3.392946377 |
| Npy2r | 3.039712243 | Myrf | -3.379653316 |
| Foxc1 | 3.037269013 | Tmem159 | -3.379510798 |
| Apoe | 3.023137061 | Ret | -3.375481099 |
| Scn10a | 3.018074335 | Cd59a | -3.367857367 |
| Tlr6 | 3.015470859 | Slc17a8 | -3.349811351 |
| Baiap2l1 | 3.011716802 | Slit2 | -3.347995862 |
| Fam43b | 3.011524897 | Apod | -3.345802811 |
| Gbp10 | 2.9906383 | Arntl2 | -3.340599725 |
| Synpo2l | 2.98887623 | Kctd19 | -3.334151869 |
| Samsn1 | 2.982233376 | Rbp3 | -3.323666317 |
| Gm1604b | 2.962660875 | Ifitm1 | -3.323664156 |
| Epx | 2.957136201 | Veph1 | -3.314188028 |
| Egr1 | 2.94532322 | 4933411K16Rik | -3.313098932 |
| Unc13c | 2.929221172 | Angpt4 | -3.313096375 |
| Krt14 | 2.921874411 | Gjc3 | -3.3039095 |
| Chrm3 | 2.910403595 | Capn11 | -3.292896716 |
| Ifi44 | 2.904876113 | Elovl3 | -3.26964341 |
| Glycam1 | 2.894862963 | Hapln2 | -3.237649121 |
| Fgf8 | 2.8908934 | Flt4 | -3.237412715 |

| | | | |
|---|---|---|---|
| Tmem71 | 2.890891022 | Ephx4 | -3.23567259 |
| Actg2 | 2.890888277 | Rab26 | -3.229114704 |
| Clec4e | 2.851771824 | Muc6 | -3.229021478 |
| Asb2 | 2.848519914 | Nfe2l3 | -3.215170174 |
| Rph3a | 2.848274692 | Mylk2 | -3.207133998 |
| Sptssb | 2.829434828 | Piezo2 | -3.202076206 |
| Fxyd3 | 2.826853824 | Flnc | -3.198930083 |
| Dbh | 2.826852417 | Foxd1 | -3.197115468 |
| Egr3 | 2.789316245 | Popdc2 | -3.185682595 |
| Ccr2 | 2.782025605 | Gata3 | -3.178420862 |
| Gna15 | 2.763975492 | Rab3b | -3.169551229 |
| Olfr1392 | 2.749659795 | Lypd6b | -3.162812728 |
| Fcgr1 | 2.743916856 | Sgk2 | -3.139728962 |
| Bspry | 2.737699477 | Tmem204 | -3.139469429 |
| Hcrtr1 | 2.731362212 | Sytl5 | -3.13862172 |
| Sycp1 | 2.729798765 | Ttc21a | -3.132409053 |
| Rasgrf1 | 2.722233844 | S1pr5 | -3.131770765 |
| Itk | 2.707267509 | Mylk3 | -3.126303602 |
| Ccl5 | 2.695346039 | Klrg1 | -3.120745167 |
| Wnt1 | 2.689369616 | Lpcat2 | -3.11982106 |
| Zpbp2 | 2.682650093 | Col12a1 | -3.11980791 |
| Hs3st4 | 2.676181324 | Esr2 | -3.113659237 |
| Kirrel2 | 2.673021238 | Thbs2 | -3.113283076 |
| Fetub | 2.671048291 | Rhox5 | -3.110128789 |
| Tacr3 | 2.665205855 | Gbp6 | -3.107478885 |
| Mettl11b | 2.652699433 | Col8a2 | -3.105792327 |
| Selplg | 2.652042588 | Mgst2 | -3.099381425 |
| Sla | 2.640397466 | Ermn | -3.09863447 |
| Lrrc25 | 2.629597422 | Cntn3 | -3.097906489 |
| Pax4 | 2.61527402 | Cdk18 | -3.08812629 |
| Dpep2 | 2.612108377 | Myo1d | -3.081809351 |
| Ncf4 | 2.612071378 | Itgb3 | -3.074815982 |
| Prss41 | 2.611974462 | Cabp4 | -3.067923462 |
| Egr2 | 2.593415631 | Treh | -3.063869448 |
| Slc6a11 | 2.586064353 | Cdh7 | -3.063863712 |
| Hp | 2.583408233 | Pm20d1 | -3.060417267 |
| Gm9992 | 2.57686017 | Myo18b | -3.058316185 |
| Fcrlb | 2.576857991 | Il12rb1 | -3.042707814 |
| Isl2 | 2.575978175 | Ahnak | -3.041502647 |
| Mrc1 | 2.553214439 | Pax9 | -3.038440032 |
| Ppp1r1b | 2.546294082 | Tm4sf1 | -3.03257173 |
| Klhl38 | 2.538076101 | Slc9a3 | -3.029859277 |
| Itgax | 2.538071814 | Atp8b1 | -3.017479885 |
| Myadml2 | 2.526543348 | 4930467E23Rik | -3.016770324 |
| Phyhip | 2.526505285 | Sbspon | -3.01484055 |
| Pnoc | 2.519624119 | Crym | -3.007657143 |
| Tnip3 | 2.496811138 | Hes3 | -3.006237329 |
| Cpne6 | 2.487721969 | Maats1 | -2.993803752 |
| Apoc1 | 2.482460867 | Plec | -2.972073523 |
| Nlrp3 | 2.482375024 | Fa2h | -2.970129893 |
| Plcb2 | 2.479950929 | Tinagl1 | -2.968898612 |
| Tnfaip8l3 | 2.45667488 | Cnbd2 | -2.955503615 |
| Acaa1b | 2.456147792 | Tspan2 | -2.950561882 |
| Epha8 | 2.437937828 | Pth2r | -2.942668803 |
| Hist1h2ad | 2.4214208 | Gucy2e | -2.935715874 |
| Gm813 | 2.420148538 | Chrna1 | -2.935713754 |
| Serpinb7 | 2.411817053 | Casq1 | -2.934365956 |
| Igfbp4 | 2.397401279 | Stx19 | -2.929921385 |

| | | | |
|---|---|---|---|
| C3 | 2.393535633 | F2rl1 | -2.918822682 |
| Rasgef1a | 2.382212459 | Itga2 | -2.918398422 |
| Mpo | 2.377621935 | Mboat1 | -2.907621089 |
| Aldh1a7 | 2.375893101 | F5 | -2.897284828 |
| Csf2rb | 2.375468267 | Ly75 | -2.897195521 |
| Apoc2 | 2.374629829 | P2rx2 | -2.891511532 |
| Prr15l | 2.367243667 | C1qtnf3 | -2.883405741 |
| Slamf1 | 2.36724042 | Sytl3 | -2.883405009 |
| Hnmt | 2.364907435 | Bpifc | -2.883402422 |
| Icam5 | 2.363923295 | Tmc5 | -2.877502418 |
| Clec4d | 2.36023286 | Fam47e | -2.869046927 |
| Runx3 | 2.359848204 | Hkdc1 | -2.864233004 |
| Ptger3 | 2.350586814 | Sspo | -2.863290158 |
| Ceacam16 | 2.330414339 | Ostn | -2.863083839 |
| Rnf43 | 2.316595405 | Abcg3 | -2.863076196 |
| Cyp4f15 | 2.299747047 | Cldn14 | -2.859105775 |
| Gstm1 | 2.296173148 | Pla2g2c | -2.854610409 |
| Gimap1 | 2.290044446 | Nxf2 | -2.854558202 |
| Tmem221 | 2.290040795 | Ankrd2 | -2.854554669 |
| Tfap2a | 2.288246245 | Abcc2 | -2.854553308 |
| Arhgap36 | 2.287024112 | Mettl21c | -2.854551741 |
| Oprm1 | 2.274267242 | Rnf183 | -2.854549972 |
| Atoh7 | 2.27426484 | Gm6583 | -2.8496882 |
| C1qtnf2 | 2.2742638 | Pla2g10 | -2.844806259 |
| Cbln4 | 2.261812238 | Cyp1a1 | -2.843184857 |
| Slc6a13 | 2.258171345 | Tnnt2 | -2.837511192 |
| Golga7b | 2.257938192 | Parvb | -2.833312582 |
| Cd300lf | 2.256000672 | Susd5 | -2.833257275 |
| Sh3tc1 | 2.247101278 | Cd80 | -2.8326462 |
| Clec5a | 2.244301921 | Col4a6 | -2.830832833 |
| Ldlrad2 | 2.244294221 | Ptk6 | -2.828878283 |
| Cd52 | 2.24065517 | 4930444G20Rik | -2.82887611 |
| Gm5741 | 2.235150397 | Mrgprf | -2.825118615 |
| Kcnc4 | 2.234758631 | Arhgef28 | -2.805673561 |
| Pld4 | 2.229844516 | Slc6a20b | -2.783930626 |
| Glp1r | 2.206246342 | Zfp951 | -2.783789528 |
| Tpd52l1 | 2.202967596 | Col16a1 | -2.781820599 |
| Cnnm1 | 2.191852062 | Ulbp1 | -2.779247644 |
| Gpr6 | 2.181434053 | Mog | -2.776958546 |
| Lrrc3b | 2.174889441 | Padi2 | -2.766445927 |
| Olfr1344 | 2.17488777 | Otog | -2.764850925 |
| Bsx | 2.174880665 | Atp10b | -2.7599242 |
| Opn5 | 2.174880483 | Nlrp1a | -2.758202492 |
| Adprhl1 | 2.174690536 | Crybb3 | -2.752999713 |
| A630001G21Rik | 2.172851279 | Dcdc2c | -2.736400346 |
| Adra2a | 2.169607846 | Tfap2c | -2.734143503 |
| Ankk1 | 2.148635843 | Stra6 | -2.73267254 |
| Cd14 | 2.129969597 | Jph2 | -2.731582256 |
| Ranbp3l | 2.127672002 | Col18a1 | -2.729552948 |
| Prok2 | 2.125879673 | Col20a1 | -2.726151176 |
| Slc27a2 | 2.12317788 | Vmn2r29 | -2.719730398 |
| F10 | 2.123039311 | Ahrr | -2.713312753 |
| Efhc2 | 2.109524266 | Clic6 | -2.713074643 |
| Scgb1b30 | 2.097683986 | Il1r1 | -2.710692907 |
| Eva1a | 2.090768055 | Lax1 | -2.709962535 |
| Cpne5 | 2.088358583 | Prelp | -2.705196221 |
| Crhr1 | 2.086137965 | Opalin | -2.704839189 |
| Lck | 2.079785612 | Unc5b | -2.70383542 |

| | | | |
|---|---|---|---|
| Srgn | 2.077624654 | Tmem132d | -2.70214402 |
| Cldn10 | 2.077289399 | Btbd16 | -2.699520166 |
| Zmat4 | 2.073845709 | Apol6 | -2.698893458 |
| Palmd | 2.066553722 | Nipal4 | -2.696521005 |
| H2-T10 | 2.066439047 | Rassf10 | -2.693612439 |
| Cbr3 | 2.064841678 | Exph5 | -2.692249462 |
| Rs1 | 2.062784343 | Otop1 | -2.69104934 |
| Lrrn4 | 2.062480636 | Klhl31 | -2.691045742 |
| Galnt9 | 2.062155051 | Sftpb | -2.688034648 |
| Pla2g4e | 2.053967799 | Vsx2 | -2.688032062 |
| Proser2 | 2.052664779 | Orm2 | -2.688031961 |
| Scel | 2.046251212 | Gpr18 | -2.688031577 |
| Adam1b | 2.045001992 | Psmb11 | -2.688031184 |
| Nos2 | 2.042508087 | Olfr543 | -2.686885337 |
| Ccr5 | 2.041639592 | Sema3g | -2.686804646 |
| Fgf5 | 2.039859848 | Barx2 | -2.685675788 |
| Fosb | 2.038199414 | Serpine1 | -2.684263335 |
| Naip5 | 2.03031144 | Sgms2 | -2.680551682 |
| Mroh7 | 2.023056056 | Styxl1 | -2.677093006 |
| Fgf9 | 2.022624459 | AW551984 | -2.676160369 |
| Nell1 | 2.020615739 | Tex26 | -2.671876686 |
| Aadat | 2.005981146 | Klk7 | -2.67187408 |
| Il21r | 2.000746585 | Arhgef5 | -2.662459514 |
| Fabp4 | 2.000745017 | Mnda | -2.662322263 |
| Me3 | 1.999290934 | Cidea | -2.66219344 |
| Kcnj4 | 1.998580941 | Morc2b | -2.659999591 |
| Cyth4 | 1.997727571 | Enpp2 | -2.654274874 |
| Klk10 | 1.98900708 | Scara3 | -2.652453353 |
| Car3 | 1.987492606 | Neil3 | -2.652079052 |
| Clec3b | 1.985454781 | Col23a1 | -2.649634541 |
| Cxcl2 | 1.9853916 | Mmp3 | -2.645779675 |
| Spock3 | 1.984539093 | Scgb3a1 | -2.643757517 |
| Wnt4 | 1.980737001 | Angptl2 | -2.634838577 |
| Baalc | 1.968173112 | Glp2r | -2.629091351 |
| Tmem255b | 1.967448713 | Medag | -2.620655995 |
| Lta | 1.964463164 | Pappa | -2.615139645 |
| Rasal3 | 1.964336329 | S100b | -2.611985467 |
| Itih4 | 1.962658703 | Pmp2 | -2.60523196 |
| Mybpc1 | 1.959312856 | Klk8 | -2.601657702 |
| Fxyd1 | 1.95499176 | Bcas1 | -2.599868263 |
| Rbp4 | 1.953112694 | Kif14 | -2.592344952 |
| Htr7 | 1.952337564 | Tnni1 | -2.590764021 |
| Cd163 | 1.952205562 | Ddc | -2.588994066 |
| Lyz2 | 1.944153155 | Mns1 | -2.587217995 |
| BC049715 | 1.94340725 | Bdkrb2 | -2.585082403 |
| Myh8 | 1.939170207 | Il1r2 | -2.579437637 |
| Rgs11 | 1.934030825 | Elovl7 | -2.576357642 |
| Wscd2 | 1.931181458 | Igbp1b | -2.576189151 |
| Smco3 | 1.929056116 | Prkg1 | -2.56250478 |
| Tbx2 | 1.928321424 | Agtr1a | -2.56064377 |
| Atp6v1e2 | 1.924694891 | Afap1l2 | -2.556225453 |
| Fxyd7 | 1.923814056 | Pof1b | -2.555734504 |
| Ifit1 | 1.923786225 | Sox7 | -2.552312404 |
| Ripk4 | 1.921866961 | Enpp3 | -2.546786811 |
| Ttpa | 1.92086168 | Cdsn | -2.546352322 |
| Nefh | 1.914478817 | Spata18 | -2.54604539 |
| Prkcg | 1.913942599 | Adamts16 | -2.54604404 |
| Adora2a | 1.910819931 | Pla2r1 | -2.545719479 |

| | | | |
|---|---|---|---|
| Klhl6 | 1.908955572 | Xirp1 | -2.54283496 |
| Slc47a2 | 1.890891636 | Rab44 | -2.541086088 |
| Pde6b | 1.890891071 | Kif19a | -2.537791865 |
| Plscr5 | 1.890889996 | Clec2d | -2.535036257 |
| Gpr132 | 1.890888883 | Heyl | -2.529969994 |
| Fam83b | 1.890886977 | Nov | -2.522674414 |
| Tmem158 | 1.890522905 | Rasgef1b | -2.522088591 |
| Itpka | 1.882428668 | Adamts4 | -2.522029462 |
| Tnfrsf14 | 1.875009495 | Vwf | -2.517800969 |
| Irgc1 | 1.875000008 | Hfm1 | -2.517686609 |
| Oprk1 | 1.874996629 | Otx2 | -2.517463074 |
| Siglec1 | 1.868506648 | Slc12a3 | -2.517313561 |
| Rbpjl | 1.868127185 | Tph2 | -2.515867931 |
| Lpo | 1.86666659 | Tmem88b | -2.51490219 |
| Pdzd3 | 1.865294764 | Tfap2e | -2.506532115 |
| Rasl11a | 1.860996841 | Asap3 | -2.504956859 |
| Gstm6 | 1.855005728 | Msx1 | -2.502201883 |
| Wnt2 | 1.840600734 | Fhl2 | -2.49727203 |
| Prmt8 | 1.836506356 | Olfr920 | -2.495679133 |
| Klf2 | 1.83599421 | Ddr2 | -2.495425543 |
| Orm1 | 1.82685391 | Car9 | -2.488124502 |
| Kcnv1 | 1.826050469 | Met | -2.484523958 |
| Sncb | 1.825219554 | Efcab1 | -2.47639853 |
| Sp100 | 1.822324603 | Mmel1 | -2.474695118 |
| Trhr | 1.819175723 | Klk11 | -2.474694672 |
| 4930502E18Rik | 1.813691087 | P2rx7 | -2.46870075 |
| Acot11 | 1.810814272 | Thbd | -2.468060014 |
| Lyz1 | 1.806224041 | Twist2 | -2.466310679 |
| Osm | 1.80486203 | Fbxl13 | -2.462591829 |
| Csf1r | 1.804762473 | Npsr1 | -2.462000319 |
| Slamf7 | 1.800660068 | Cd300lg | -2.461999915 |
| Ntsr2 | 1.791456082 | Gm6592 | -2.461999914 |
| Gstt1 | 1.790613992 | 1700034J05Rik | -2.461999284 |
| Cdh17 | 1.787740889 | St18 | -2.458659014 |
| Rbfox3 | 1.787381531 | Ifnlr1 | -2.455613591 |
| Vav1 | 1.785924963 | Opcml | -2.454156853 |
| Rasal1 | 1.77895843 | Fam216b | -2.453364603 |
| Mmp9 | 1.777254959 | Gpa33 | -2.451400755 |
| Slc25a18 | 1.772624655 | Fgf7 | -2.451031836 |
| Pla2g5 | 1.770303426 | Car10 | -2.449385593 |
| Hcls1 | 1.766016367 | Mylk4 | -2.44919867 |
| Camk2n1 | 1.760265449 | Guca2a | -2.449197173 |
| Fgf3 | 1.759850176 | Mansc4 | -2.449162796 |
| Nr5a1 | 1.759849405 | Trpv4 | -2.442582257 |
| Cd4 | 1.759845381 | Sulf1 | -2.442309983 |
| Barx1 | 1.759840763 | Hhip | -2.441791364 |
| Gm5105 | 1.754625354 | Chst8 | -2.441041568 |
| Oaf | 1.749693923 | Gimap8 | -2.436275876 |
| Wfdc6a | 1.7496565 | Prss45 | -2.436273914 |
| Npw | 1.749655793 | Olfr279 | -2.436270028 |
| Nkx1-2 | 1.749655205 | Mki67 | -2.436192099 |
| Neurog2 | 1.747313824 | Ror1 | -2.430831962 |
| Kcnk12 | 1.746827597 | Ooep | -2.429766644 |
| Mmp12 | 1.746638944 | Tmem74 | -2.428967738 |
| Epas1 | 1.746101131 | Sntg1 | -2.427594591 |
| Cds1 | 1.744015457 | Gsc | -2.426573447 |
| Bin2 | 1.742851263 | Naalad2 | -2.426012333 |
| Dhrs2 | 1.736869372 | Trim12a | -2.421021002 |

| | | | |
|---|---|---|---|
| Arc | 1.73537535 | Eid3 | -2.419471434 |
| C1ra | 1.733276042 | Ngf | -2.417588565 |
| Dennd2d | 1.730480439 | Igfbp6 | -2.417240275 |
| Efna3 | 1.726022409 | Olfr545 | -2.416672576 |
| Mchr1 | 1.722759727 | Armc3 | -2.415078994 |
| Sidt1 | 1.720537806 | Igf2 | -2.413261658 |
| Hoxc9 | 1.716541228 | Igf2bp1 | -2.412715438 |
| Itgam | 1.714022729 | Loxl2 | -2.412347927 |
| Usp44 | 1.700965214 | Ldb3 | -2.412162196 |
| Ms4a6b | 1.700613116 | Adora3 | -2.410077278 |
| Ccl22 | 1.697104777 | Itga11 | -2.40395764 |
| Lrmp | 1.689992887 | Tek | -2.401859093 |
| Folh1 | 1.688210791 | Dusp26 | -2.401222646 |
| Ctss | 1.687420652 | Lhx9 | -2.398789583 |
| BC024139 | 1.685819551 | Ddx4 | -2.398647497 |
| Pgr | 1.685252372 | Ngfr | -2.387633155 |
| Tifa | 1.683797199 | Slc2a12 | -2.380378707 |
| Gda | 1.682347828 | Slc10a6 | -2.377894442 |
| Ccl9 | 1.660884585 | Cox4i2 | -2.375280046 |
| Tlcd1 | 1.659541884 | Nfasc | -2.37441999 |
| Cacna1i | 1.659367821 | Atp8b3 | -2.374289063 |
| Sp7 | 1.655026667 | Nek11 | -2.373731169 |
| Caln1 | 1.65433727 | Syt15 | -2.370866173 |
| Gpr31b | 1.650055819 | Mlkl | -2.370104398 |
| Gas2l2 | 1.648336863 | Mgp | -2.366998462 |
| Plcd4 | 1.640396412 | Slc9b1 | -2.362616423 |
| Cst3 | 1.638959088 | Cyp26b1 | -2.357203422 |
| Aoah | 1.638896467 | Bmp6 | -2.353524602 |
| Ccl4 | 1.63738143 | Aldh8a1 | -2.348018866 |
| Trim75 | 1.637379221 | Grid2 | -2.346248711 |
| Inpp5j | 1.630278618 | Aspm | -2.343554644 |
| 9930111J21Rik1 | 1.63006511 | Morn5 | -2.343367131 |
| Cdkl1 | 1.629356295 | Ndrg1 | -2.337389455 |
| Ncf1 | 1.627973057 | Perp | -2.335578745 |
| Nxnl1 | 1.626001988 | St14 | -2.333654558 |
| Gpr12 | 1.624559107 | Cd244 | -2.333201225 |
| Phkg1 | 1.624262735 | Clrn1 | -2.328735557 |
| Gbp5 | 1.613923113 | Timp1 | -2.327491379 |
| Fam159a | 1.613576826 | Rho | -2.322063996 |
| Actc1 | 1.613168092 | Msln | -2.320153816 |
| Ly9 | 1.605553898 | Gsta2 | -2.320152352 |
| Syt12 | 1.605231208 | Cped1 | -2.319579994 |
| Glo1 | 1.604700314 | Tnxb | -2.31697889 |
| Cytip | 1.601660311 | Fgf18 | -2.315382009 |
| Lars2 | 1.59939962 | Clhc1 | -2.309474552 |
| Trim29 | 1.599311991 | Prune2 | -2.307741182 |
| Ripply2 | 1.597956019 | Pxdc1 | -2.305770346 |
| Baiap2l2 | 1.595228421 | Ypel2 | -2.30213091 |
| Ptpn18 | 1.588075988 | Syne2 | -2.301134484 |
| A830018L16Rik | 1.579582688 | Il13ra1 | -2.297451569 |
| Sowahb | 1.576861804 | Clmn | -2.294935197 |
| Fam167b | 1.576858413 | Tmem40 | -2.287635832 |
| S100a8 | 1.576857895 | Ddo | -2.280388215 |
| Apol9b | 1.576856122 | Lrrc9 | -2.276624366 |
| Vmn1r58 | 1.576159574 | Trh | -2.275724677 |
| Zfp345 | 1.575499022 | Hist1h3b | -2.274248829 |
| Scn7a | 1.575496105 | Sstr5 | -2.269646519 |
| Clec4a1 | 1.572052879 | Lhfpl5 | -2.269646223 |

| | | | |
|---|---|---|---|
| Rnf151 | 1.571767223 | Iyd | -2.269643757 |
| Apobec1 | 1.570703901 | Cdh12 | -2.266087307 |
| Sec14l3 | 1.568964594 | Ptpre | -2.263252821 |
| Dll3 | 1.568476853 | Rhpn2 | -2.261980917 |
| Ism1 | 1.56795832 | Pmaip1 | -2.259960326 |
| Abi3 | 1.559807241 | Ptchd4 | -2.259194051 |
| Nab2 | 1.559149507 | Clnk | -2.256839541 |
| Lypd6 | 1.557355464 | AU018091 | -2.256839231 |
| Npas4 | 1.553890457 | Angpt2 | -2.254988435 |
| 2010300C02Rik | 1.551034956 | Tm4sf4 | -2.243919082 |
| Rd3 | 1.551001103 | Pkp1 | -2.243917594 |
| Agmo | 1.542525617 | 1700047I17Rik2 | -2.243914866 |
| Bdh2 | 1.540151719 | Hsd3b3 | -2.24391393 |
| Tmco5 | 1.538073646 | Osr1 | -2.242483839 |
| Slc47a1 | 1.53807175 | Klhl4 | -2.24171678 |
| Sp110 | 1.537996453 | Mcam | -2.23602914 |
| Myo1g | 1.537878665 | Clgn | -2.234759447 |
| Lrrc6 | 1.537739679 | Uchl4 | -2.227785365 |
| C1rl | 1.53739017 | Adrb3 | -2.227784219 |
| Cabp7 | 1.537388695 | Gpr35 | -2.227600577 |
| Gpr61 | 1.534051234 | Has1 | -2.221902542 |
| Syndig1 | 1.530387899 | Lrtm1 | -2.221877011 |
| Ckmt1 | 1.524764631 | Hdc | -2.22076784 |
| Cwc22 | 1.524011943 | Tec | -2.220208527 |
| Fmo1 | 1.522649623 | Hgf | -2.219206827 |
| Csf2rb2 | 1.520373832 | Mrgprb5 | -2.217723014 |
| Ung | 1.519498213 | C7 | -2.211194639 |
| Tnf | 1.518569986 | Ptx3 | -2.210713725 |
| Ccdc85a | 1.513521874 | Bpifb5 | -2.209135519 |
| Flywch2 | 1.513127417 | Gsc2 | -2.209129956 |
| Jph4 | 1.51223961 | Tfap2b | -2.209129914 |
| Cpne4 | 1.511297141 | Abhd14b | -2.208869022 |
| Kcne1l | 1.509985777 | Prom2 | -2.206634789 |
| Npas2 | 1.50994897 | Sertm1 | -2.205098963 |
| Camk2b | 1.509635942 | Rnasel | -2.204095997 |
| Cdk5r2 | 1.508936037 | Nr1i2 | -2.201517038 |
| Il11ra1 | 1.505736211 | Pla1a | -2.201516187 |
| Ybx2 | 1.50418382 | Matn1 | -2.201367276 |
| Nfam1 | 1.500662347 | Tnc | -2.201354196 |
| Pcp4 | 1.500129212 | Slc34a3 | -2.198963491 |
| Ifitm6 | 1.499657059 | Plcg2 | -2.198264119 |
| Gstm3 | 1.499655631 | Stk32b | -2.195554361 |
| T | 1.499654133 | Cenpe | -2.195079195 |
| Galnt18 | 1.498593074 | Entpd4 | -2.194641604 |
| Sfrp5 | 1.492046641 | 4931429L15Rik | -2.18615992 |
| Plbd1 | 1.491795482 | Sema5a | -2.185964793 |
| Pak6 | 1.488237093 | Tsx | -2.183588465 |
| Nrarp | 1.4874017 | Mal | -2.182229662 |
| Dleu7 | 1.487195587 | Trim36 | -2.181335418 |
| Hddc3 | 1.48523812 | Ccdc114 | -2.181068546 |
| Paqr7 | 1.484848187 | Rfpl4 | -2.178425628 |
| Mesp2 | 1.481951896 | Gckr | -2.178424447 |
| Doc2b | 1.481209204 | Neurog1 | -2.1784215 |
| Vstm2l | 1.481153445 | Ccdc158 | -2.178420047 |
| Prrt1 | 1.477187455 | C130026I21Rik | -2.178419029 |
| Fam163b | 1.476903821 | Pamr1 | -2.177200457 |
| Ntsr1 | 1.476383427 | Tppp | -2.17439248 |
| Phospho1 | 1.473258592 | Fam187a | -2.170643543 |

| | | | |
|---|---|---|---|
| Fmnl1 | 1.467193706 | Arsj | -2.164956477 |
| Ms4a6d | 1.466914607 | Crabp2 | -2.162807695 |
| Slc5a9 | 1.466089227 | Ceacam1 | -2.162667997 |
| Rac2 | 1.463684085 | Mob3b | -2.161066126 |
| Rlbp1 | 1.458334957 | Dkk3 | -2.160323344 |
| Cd38 | 1.45420244 | Kif11 | -2.159609771 |
| Th | 1.45180168 | Prokr1 | -2.1580233 |
| Hist2h2ac | 1.451258358 | Col17a1 | -2.157843086 |
| Ajap1 | 1.450715876 | 1810024B03Rik | -2.157839231 |
| Fcgr4 | 1.445001762 | Ctse | -2.157838799 |
| Col4a4 | 1.442755214 | Cel | -2.155655291 |
| Tox2 | 1.440221662 | Myof | -2.154431061 |
| Neurod6 | 1.436736174 | Fn1 | -2.15261974 |
| Nlrp10 | 1.436735948 | Grm6 | -2.147046471 |
| Cdhr1 | 1.436659106 | Tgm1 | -2.143580457 |
| Tmem72 | 1.434245029 | Gpr152 | -2.141792551 |
| Cdhr3 | 1.433318434 | Dydc2 | -2.141789801 |
| Kremen2 | 1.431680927 | Adamtsl4 | -2.141533139 |
| Tnnt3 | 1.428706738 | E030018B13Rik | -2.137048841 |
| Foxh1 | 1.426377922 | Plxnb3 | -2.133965449 |
| Slc24a4 | 1.426218048 | Nfatc4 | -2.131835844 |
| Samd1 | 1.425227188 | Cenpf | -2.131263749 |
| Rasgrp2 | 1.423514253 | Esam | -2.130338291 |
| Ccnb1ip1 | 1.419613462 | Ccdc150 | -2.128976948 |
| Agtr2 | 1.41961107 | Myl2 | -2.127794597 |
| Sp6 | 1.418788477 | Capn3 | -2.12172048 |
| Gfra4 | 1.415683694 | Itpr2 | -2.121243506 |
| Rhcg | 1.412756477 | Col2a1 | -2.120784183 |
| Lyn | 1.409504227 | Olfr1393 | -2.120742152 |
| Slc12a5 | 1.409233601 | Eps8l2 | -2.118685588 |
| Crip1 | 1.409022276 | Gm7173 | -2.113923727 |
| Slco1a4 | 1.406091277 | Adssl1 | -2.113896972 |
| Was | 1.406091042 | 4930590J08Rik | -2.113656982 |
| 5830473C10Rik | 1.405672688 | Tssk3 | -2.113656389 |
| Acsbg1 | 1.405575749 | Pparg | -2.107107786 |
| Rasl10b | 1.405520555 | Klf17 | -2.106540445 |
| Gls2 | 1.40305438 | Trim43a | -2.106153868 |
| Hmox1 | 1.400613971 | A3galt2 | -2.105919035 |
| Psd4 | 1.399148052 | Abca13 | -2.103070368 |
| Acaa2 | 1.399077292 | Iqgap3 | -2.101289616 |
| Bend6 | 1.399031345 | Hist1h4n | -2.096824799 |
| H2-Ke6 | 1.397670721 | St6galnac2 | -2.095598109 |
| Stc2 | 1.397214663 | Gpr37 | -2.094856922 |
| Cnr2 | 1.396537014 | Itga3 | -2.094507432 |
| Mlxipl | 1.392686836 | Pde1c | -2.093596126 |
| Gdf10 | 1.392686809 | Tnni3 | -2.092127241 |
| Dmkn | 1.389825303 | Map3k6 | -2.091019462 |
| Gpr22 | 1.388933693 | Riiad1 | -2.088741414 |
| Hpdl | 1.387847274 | Itgb7 | -2.087764931 |
| Sv2c | 1.387591467 | Slc7a14 | -2.08752095 |
| Slfn2 | 1.386362149 | Mdh1b | -2.086777317 |
| Foxd2 | 1.383573659 | Pkd2l1 | -2.079040319 |
| Lrfn5 | 1.38319776 | Chrna5 | -2.076883393 |
| Gabra5 | 1.376890823 | Ptprd | -2.070889903 |
| Scn1b | 1.376278823 | Itgb4 | -2.069486819 |
| Hif3a | 1.376065355 | Fbxo32 | -2.065908119 |
| Vsig8 | 1.374454104 | Dsc2 | -2.063438275 |
| Satb2 | 1.37182617 | Plk5 | -2.063434344 |

| | | | |
|---|---|---|---|
| Fgf1 | 1.371780402 | Ifitm10 | -2.059876756 |
| Chst1 | 1.370851302 | Bgn | -2.059079757 |
| Fcrls | 1.36924072 | Zbtb42 | -2.058541059 |
| Fcer1g | 1.368810954 | Endou | -2.055439933 |
| Wnt16 | 1.367244826 | Sall4 | -2.055437914 |
| Lgals7 | 1.367243567 | Wfdc2 | -2.054975537 |
| Ckmt2 | 1.367242918 | Grem1 | -2.052737039 |
| Lgals12 | 1.367242217 | Gpr165 | -2.051197008 |
| Cdhr5 | 1.367241579 | Col9a1 | -2.050298672 |
| Otof | 1.367240412 | Arhgef19 | -2.047805971 |
| Defb42 | 1.367239362 | Fbln5 | -2.045091178 |
| Trim63 | 1.367239155 | Igsf9b | -2.043677369 |
| H2-Aa | 1.36723852 | Ugt8a | -2.040583499 |
| 4933409G03Rik | 1.367238307 | Ank3 | -2.037285804 |
| Bmx | 1.367238137 | Gm5868 | -2.036059409 |
| Hoxa7 | 1.367237866 | Kif27 | -2.035742573 |
| Ngp | 1.367237554 | Scn5a | -2.033126043 |
| Metrnl | 1.365962146 | Slc23a3 | -2.031886477 |
| Syt5 | 1.36531996 | Speer3 | -2.03188522 |
| Pde4a | 1.365308889 | Cnga3 | -2.029863008 |
| Gbp3 | 1.362823057 | Slc17a6 | -2.029860588 |
| Dlk2 | 1.358243809 | Btbd8 | -2.026079455 |
| Ryr3 | 1.358206151 | Pif1 | -2.025814457 |
| Asb5 | 1.356765618 | Clec2g | -2.0256041 |
| Fxyd5 | 1.356178835 | Dpcr1 | -2.024348947 |
| Drd4 | 1.356101668 | Arsi | -2.02076956 |
| Rhoq | 1.355995795 | Trim72 | -2.019741688 |
| Cebpa | 1.355329099 | Rinl | -2.019217386 |
| Hist1h2ah | 1.355203179 | Sox18 | -2.016775655 |
| Nupr1l | 1.352940124 | Tph1 | -2.016774865 |
| Kcnk3 | 1.352351512 | Fsd2 | -2.01677263 |
| Dio3 | 1.351667946 | Htr1f | -2.012554446 |
| Alpk3 | 1.350711257 | B3galt5 | -2.008698101 |
| Gfpt2 | 1.350356087 | Smim6 | -2.008146466 |
| Gck | 1.3495651 | Ass1 | -2.00622182 |
| Gulp1 | 1.34781762 | Cnpy1 | -2.00545526 |
| Fbxo2 | 1.346733198 | Neu4 | -2.005107503 |
| Tmem37 | 1.344158526 | Gm128 | -2.004546687 |
| Mdga1 | 1.341631742 | Aldh3a1 | -2.001692474 |
| 1700023F06Rik | 1.341268542 | Gnb3 | -2.001502248 |
| Sox9 | 1.340884758 | Amdhd1 | -2.001497494 |
| Ttc34 | 1.339338896 | Nsun7 | -2.000654456 |
| Dnm1 | 1.338909602 | Entpd2 | -1.999953388 |
| Clybl | 1.338591537 | Erbb4 | -1.997303872 |
| Mt1 | 1.33517806 | Gm6904 | -1.993803588 |
| Spta1 | 1.334772721 | Bfsp2 | -1.992616447 |
| Myh7 | 1.334007174 | Kif15 | -1.98916442 |
| Mtfp1 | 1.333991617 | Tmem151a | -1.988946693 |
| Ctf1 | 1.332211159 | Sfrp4 | -1.987689436 |
| Xlr4b | 1.331543924 | Kbtbd12 | -1.9860639 |
| Cxcl12 | 1.330643944 | Bex6 | -1.986063528 |
| Cyp27b1 | 1.328967129 | 1600014C23Rik | -1.986061794 |
| Tgm3 | 1.326633047 | Gsdma | -1.986060456 |
| Ptges2 | 1.32449076 | Zdhhc22 | -1.982651981 |
| Wif1 | 1.323249615 | Vcan | -1.982533616 |
| Pcsk5 | 1.322356161 | Tmem116 | -1.979690154 |
| Sypl2 | 1.322081379 | Slc45a3 | -1.975895607 |
| Prr18 | 1.321870833 | Pde9a | -1.97520147 |

| | | | |
|---|---|---|---|
| Nrtn | 1.317885922 | D7Ertd443e | -1.974024611 |
| Klf10 | 1.316083338 | Ptgr1 | -1.972617345 |
| Rgl3 | 1.315301764 | Bcl2l15 | -1.972022737 |
| Ndrg2 | 1.307143767 | Il4 | -1.97046556 |
| Upp1 | 1.304609845 | Reln | -1.967235443 |
| Kcnh1 | 1.302562113 | Tspan17 | -1.962986558 |
| Mgst3 | 1.301573935 | Sh2d4b | -1.962668299 |
| Wtip | 1.300883077 | Fam163a | -1.955736252 |
| 1110065P20Rik | 1.300857275 | Emp1 | -1.955120457 |
| Arhgap25 | 1.29442486 | Loxl1 | -1.954656243 |
| Inha | 1.293920199 | Nlrc5 | -1.953094508 |
| Ascl2 | 1.29296259 | Stmn4 | -1.948641099 |
| Cebpb | 1.292360641 | Ncam2 | -1.947796132 |
| Fam151a | 1.29222034 | Scn9a | -1.947582452 |
| Krt80 | 1.290045998 | Adamts2 | -1.944646844 |
| Klrk1 | 1.290042832 | Dnajb7 | -1.941569626 |
| Catsper3 | 1.290042784 | Cnn1 | -1.941039375 |
| Nme8 | 1.290042549 | Pyroxd2 | -1.939055595 |
| Gabrp | 1.290042338 | Ahr | -1.936249472 |
| Igfn1 | 1.290040751 | Pdlim2 | -1.935952913 |
| Gm5111 | 1.290040592 | Stk32a | -1.934571794 |
| Cdcp1 | 1.290040241 | Adgb | -1.9304823 |
| Tsga13 | 1.29004012 | Mx2 | -1.930111402 |
| Dpp4 | 1.290038803 | Folr2 | -1.927818291 |
| Fam83a | 1.290038682 | Btc | -1.927093451 |
| 1700125H20Rik | 1.290038483 | Galnt12 | -1.926625422 |
| Ikzf3 | 1.290036643 | Arhgap11a | -1.926359266 |
| Psg28 | 1.290035473 | Irx1 | -1.92569898 |
| Mecom | 1.28723958 | Kif4 | -1.923686246 |
| Ptgis | 1.286542016 | Esyt3 | -1.917583164 |
| Srm | 1.284647291 | Capn5 | -1.915448511 |
| Kcnk1 | 1.28341516 | S100a10 | -1.914951068 |
| Arl5c | 1.281638996 | Tbx3 | -1.913840517 |
| Tlr4 | 1.280531983 | Chrna3 | -1.913081601 |
| Pnp | 1.280151631 | Gnrh1 | -1.903843121 |
| Abhd16b | 1.279733762 | Gapt | -1.902321771 |
| Ngef | 1.278721475 | Tbx19 | -1.902315057 |
| Agap2 | 1.276689606 | Ccr4 | -1.902314163 |
| Il10ra | 1.276215174 | Sucnr1 | -1.898484387 |
| Cpvl | 1.274268762 | Gap43 | -1.892477786 |
| Rpl39l | 1.274266177 | Zfp872 | -1.891850944 |
| 1810011H11Rik | 1.274263661 | Plxna4 | -1.8912937 |
| Parvg | 1.273756715 | Cd82 | -1.888215862 |
| 1700001O22Rik | 1.272621143 | Hrasls | -1.885058043 |
| Lepr | 1.270229058 | Grpr | -1.883402841 |
| Fbll1 | 1.265224996 | Olfr1361 | -1.883400127 |
| Bcl2a1b | 1.264527309 | Pla2g4f | -1.883398239 |
| Prg4 | 1.263321319 | Hao1 | -1.883396659 |
| Tceal7 | 1.263217265 | Hsd11b2 | -1.883292963 |
| Eno2 | 1.262718541 | Adam5 | -1.883246597 |
| Ablim3 | 1.259661525 | Il4ra | -1.88006219 |
| Slpi | 1.257799816 | Mtfr2 | -1.87894722 |
| Nlrp6 | 1.254576638 | Hist1h2bm | -1.877037075 |
| Trpc7 | 1.252457788 | Cdca2 | -1.876672908 |
| Ltc4s | 1.250092827 | Col5a1 | -1.875522985 |
| Cort | 1.247102583 | Kif6 | -1.872157451 |
| Cd37 | 1.246604362 | Abca6 | -1.87065008 |
| Itih3 | 1.243381739 | Stx11 | -1.870583951 |

| | | | |
|---|---|---|---|
| Zic3 | 1.243264841 | Atoh8 | -1.869845185 |
| Hist1h2bp | 1.242960616 | Zfp488 | -1.866954988 |
| Rerg | 1.24127818 | Rgs9 | -1.866226337 |
| Nog | 1.239963405 | Iapp | -1.866212899 |
| Tmem82 | 1.238710922 | Efna1 | -1.866006056 |
| Wdr72 | 1.238259264 | Pik3cd | -1.865535074 |
| Ptk2b | 1.237251566 | Hand1 | -1.864235624 |
| Rnd1 | 1.232261754 | Acox2 | -1.864234457 |
| Dkk1 | 1.231261356 | Sis | -1.864232841 |
| Caly | 1.228446199 | 4933403O08Rik | -1.864232772 |
| Cox6a2 | 1.225429052 | Gm15217 | -1.864231793 |
| Mboat4 | 1.225428965 | Il13 | -1.864230486 |
| Apoa1 | 1.225424277 | Cxcl9 | -1.864229332 |
| Comtd1 | 1.2248908 | Gp5 | -1.863733541 |
| Raver2 | 1.222667771 | Tac4 | -1.863433966 |
| Ppp1r14b | 1.2222665 | Smim5 | -1.863427946 |
| Gpd1 | 1.219446606 | Rnaset2b | -1.862932154 |
| Erf | 1.218243088 | Cpm | -1.862826586 |
| Kcns1 | 1.218130837 | Mak | -1.857544049 |
| Mctp1 | 1.21668618 | Dlgap5 | -1.853931847 |
| Cnih3 | 1.215178455 | Csgalnact1 | -1.853044895 |
| Rom1 | 1.213391189 | Srpk3 | -1.851410178 |
| Psd | 1.213046979 | Tal2 | -1.850042283 |
| 1700029I15Rik | 1.212887227 | Dock9 | -1.849513525 |
| Aldoc | 1.211812197 | Btn1a1 | -1.846081827 |
| Rfx4 | 1.211678049 | Fcgbp | -1.846081237 |
| Lxn | 1.211353611 | Odf4 | -1.846080738 |
| Lor | 1.209403093 | Ggt6 | -1.844802771 |
| Trpc6 | 1.206580911 | Kif23 | -1.844483848 |
| Arg2 | 1.206301281 | Tenm1 | -1.842693955 |
| Rhbdl1 | 1.205749482 | Fat4 | -1.842143697 |
| Hist1h2bk | 1.203959414 | Man1c1 | -1.838273564 |
| Klf16 | 1.19956902 | Spn | -1.837510423 |
| Cftr | 1.198395876 | Cst7 | -1.837506653 |
| Polr2f | 1.197528596 | Slc37a2 | -1.83675372 |
| Il1a | 1.197370802 | Nuf2 | -1.835916123 |
| Vnn3 | 1.197068153 | Tlr5 | -1.834994821 |
| Gm826 | 1.197066324 | Clcf1 | -1.834112923 |
| Meox1 | 1.197066225 | Cep128 | -1.833613281 |
| Junb | 1.195691403 | Slc41a3 | -1.83123793 |
| Inpp5d | 1.193689382 | Ldlrap1 | -1.829145672 |
| Lalba | 1.189972833 | Tmem63a | -1.825222995 |
| Cpa6 | 1.189971603 | Mtnr1a | -1.825118678 |
| Ptprcap | 1.187061454 | Bik | -1.825111749 |
| Dusp6 | 1.18705629 | Batf | -1.825109995 |
| Pik3c2g | 1.186316475 | Ace | -1.822905737 |
| Lrit1 | 1.18631432 | Col6a1 | -1.821603321 |
| Gipc3 | 1.186313785 | Echdc3 | -1.820201558 |
| C1qtnf4 | 1.184632038 | Itga7 | -1.819657484 |
| Nr2f6 | 1.184067868 | Abca5 | -1.817488137 |
| Pdia2 | 1.18258397 | Sag | -1.817420867 |
| Rtp1 | 1.182359234 | Postn | -1.817385747 |
| 2610528A11Rik | 1.181344002 | Aldh1l2 | -1.816840149 |
| Cyp2s1 | 1.178059555 | Hgfac | -1.816392317 |
| Ascl4 | 1.177727786 | Layn | -1.815265604 |
| 2310039H08Rik | 1.177504254 | Serpinf1 | -1.81469698 |
| Cygb | 1.17656956 | Krt73 | -1.812481214 |
| Upb1 | 1.174889495 | Slitrk6 | -1.810510935 |

| | | | |
|---|---|---|---|
| Lmx1b | 1.174887675 | Anxa1 | -1.807525471 |
| C8a | 1.17488599 | Pvr | -1.807483486 |
| Tgtp1 | 1.174885966 | Lct | -1.807159476 |
| Cyp1a2 | 1.174885058 | Capn6 | -1.8069022 |
| Fgf16 | 1.174884174 | Plekha6 | -1.805644511 |
| C1ql4 | 1.17488375 | Dst | -1.802011339 |
| Gm8909 | 1.174883292 | Ermap | -1.801153341 |
| Mroh5 | 1.174693487 | Arhgef39 | -1.800596528 |
| Glul | 1.172337497 | Ces5a | -1.797220944 |
| Prr22 | 1.171605824 | Plscr4 | -1.796363955 |
| Kcng2 | 1.168564325 | Itih5 | -1.795246633 |
| Fpr1 | 1.167017599 | Pex5l | -1.793374592 |
| Akap3 | 1.167008917 | Lrriq1 | -1.789964443 |
| Golph3 | 1.165148358 | Prr19 | -1.789059502 |
| Cnih2 | 1.161764942 | Nid1 | -1.787969321 |
| Asic4 | 1.161693006 | Sdsl | -1.787757565 |
| Vstm2a | 1.160660171 | 4933408B17Rik | -1.786487895 |
| Tlr2 | 1.160214331 | Arhgap6 | -1.785650395 |
| Pde1b | 1.159946644 | Upp2 | -1.784019941 |
| Aanat | 1.157234194 | Tubb4a | -1.782424397 |
| Phgdh | 1.156475589 | Mroh2b | -1.781529356 |
| Fam174b | 1.15451938 | Cryl1 | -1.779515767 |
| Lyplal1 | 1.154291735 | Gdf6 | -1.77812104 |
| Chchd7 | 1.154055411 | Qrfp | -1.776900493 |
| B3gnt3 | 1.153924094 | Snx30 | -1.776573115 |
| Foxe1 | 1.148638328 | Sec14l5 | -1.776075968 |
| Prss56 | 1.148229216 | Cldn20 | -1.775793399 |
| 1110032F04Rik | 1.14671658 | Adra2b | -1.7744651 |
| C1qb | 1.1464117 | Dock5 | -1.773304384 |
| Tmem238 | 1.145615901 | Gabra3 | -1.772561127 |
| Kcnj16 | 1.144888243 | Ifih1 | -1.770285248 |
| Snta1 | 1.14462141 | Ankle1 | -1.768571581 |
| Arhgdib | 1.144260859 | P2ry2 | -1.767864626 |
| Gm14295 | 1.143308377 | Cybrd1 | -1.767548008 |
| Scg2 | 1.142773793 | Syngr4 | -1.767261549 |
| Espn | 1.142387346 | Hist1h4b | -1.764446483 |
| C2 | 1.14171869 | Ltbp1 | -1.763028814 |
| Kank3 | 1.140952789 | Ppp1r3b | -1.76296588 |
| Pus7l | 1.138384879 | Kcna6 | -1.762440605 |
| Cacna1e | 1.137789274 | Efemp1 | -1.760317129 |
| Crybb1 | 1.136492882 | Rbms1 | -1.75978034 |
| Rbp7 | 1.13576504 | Car6 | -1.759140524 |
| Cilp2 | 1.133552759 | Cntn2 | -1.758524688 |
| Rapgefl1 | 1.132168038 | Ptprt | -1.758336272 |
| Klhdc8b | 1.129723024 | Lrp2 | -1.757811671 |
| Mical1 | 1.127230822 | Smad6 | -1.755324664 |
| Il7 | 1.125863934 | Mss51 | -1.754104764 |
| Rrp9 | 1.123587495 | Itga1 | -1.753498877 |
| Hist2h3c1 | 1.123147039 | Mapk13 | -1.753356538 |
| Stpg2 | 1.122962987 | Igfbp5 | -1.753086434 |
| Ppara | 1.122265364 | Spef2 | -1.752359612 |
| Stxbp6 | 1.12036109 | Fank1 | -1.749624014 |
| Spock1 | 1.12010401 | Glb1l2 | -1.747291976 |
| Ltb4r2 | 1.120065969 | Lrguk | -1.745852958 |
| Aass | 1.119165482 | Syt4 | -1.745168812 |
| Il1rn | 1.118646843 | Tiam2 | -1.742320863 |
| Tcerg1l | 1.117246857 | Cacna2d2 | -1.739673415 |
| Mfsd2a | 1.113868826 | Pmch | -1.739193183 |

| | | | |
|---|---|---|---|
| Traf4 | 1.113388796 | Igsf10 | -1.737658204 |
| Zbtb7c | 1.113074196 | Trim43c | -1.735894795 |
| Il15ra | 1.110692538 | Il11 | -1.735791513 |
| Fxyd2 | 1.110635475 | Crispld1 | -1.735190106 |
| Trmt61a | 1.109358405 | Ddx60 | -1.733192592 |
| Mrpl12 | 1.107451576 | Itgae | -1.731044154 |
| Gm5617 | 1.106720915 | Styk1 | -1.728897941 |
| Pip5k1b | 1.105605339 | Otoa | -1.727228636 |
| Rit2 | 1.105517435 | Col6a2 | -1.727030343 |
| Trem2 | 1.104083784 | Adam32 | -1.726743238 |
| Pdzd2 | 1.102754941 | Col6a3 | -1.725770288 |
| Zfp456 | 1.1018819 | Cd44 | -1.725349089 |
| Wipf3 | 1.101663617 | Fzd6 | -1.719980591 |
| Gjd2 | 1.100968574 | 4930562C15Rik | -1.719383863 |
| Pp2d1 | 1.10087623 | Sh3pxd2a | -1.716607738 |
| Gm21949 | 1.10009815 | Scd3 | -1.716048286 |
| Lcp2 | 1.099456551 | 1190007I07Rik | -1.715152462 |
| Adam33 | 1.097686941 | Colec11 | -1.712780285 |
| Tmprss11g | 1.097686 | Pcsk9 | -1.712472979 |
| Gdf3 | 1.097684561 | B430306N03Rik | -1.709960739 |
| Prdm12 | 1.097683583 | Tcf23 | -1.709959824 |
| Zbbx | 1.097681161 | Rad51ap2 | -1.709959259 |
| Card10 | 1.097071677 | Olfr1318 | -1.709958264 |
| Tusc1 | 1.096588048 | Ffar3 | -1.709956693 |
| Olfml1 | 1.096379732 | Olfr95 | -1.709953742 |
| Cntnap5a | 1.096109368 | Tead4 | -1.708155591 |
| Adprhl2 | 1.095868956 | Kank2 | -1.704291984 |
| Pdzd9 | 1.094282175 | Mlf1 | -1.703905143 |
| Zcchc12 | 1.091374858 | Map1b | -1.700692368 |
| Trim65 | 1.09117345 | Bmp8b | -1.700535773 |
| Dusp2 | 1.090779525 | Chodl | -1.697490262 |
| Ccl2 | 1.089986916 | Ubc | -1.692132394 |
| Gm2a | 1.089632695 | Rorc | -1.691927578 |
| Scube1 | 1.08830953 | Ctsk | -1.691881859 |
| Hapln4 | 1.087321526 | Vwc2 | -1.691462377 |
| Rln1 | 1.086118365 | Enpep | -1.691045802 |
| Npy1r | 1.085558519 | Ecscr | -1.691043982 |
| Tcp10c | 1.08308805 | Pax3 | -1.691041405 |
| Cited1 | 1.079978607 | Cyp4b1 | -1.691040279 |
| Tyrobp | 1.079510275 | Abcc9 | -1.691037311 |
| Jakmip1 | 1.078998346 | Tdh | -1.688031576 |
| Pdk4 | 1.078991364 | Tmem217 | -1.688027539 |
| Laptm5 | 1.077348344 | Prph | -1.684742684 |
| Zdbf2 | 1.076983066 | 3632451O06Rik | -1.684378789 |
| Olfm3 | 1.076715523 | Plin1 | -1.681491532 |
| Tdrd9 | 1.075586803 | Cit | -1.681386519 |
| Prdx6 | 1.075289387 | Vmn1r32 | -1.680749331 |
| Dhh | 1.075229998 | S1pr3 | -1.679624871 |
| Tmem121 | 1.074643027 | Morn3 | -1.679440964 |
| E030030I06Rik | 1.074480054 | Vwa2 | -1.677090916 |
| Tmem240 | 1.074105852 | Tnr | -1.675118831 |
| Ubiad1 | 1.073610773 | Mmp2 | -1.67390703 |
| Grin1 | 1.072358541 | Macc1 | -1.673799205 |
| Ccdc69 | 1.071738222 | Grm7 | -1.673229071 |
| Itgb2 | 1.06951795 | Rhbdl2 | -1.67290435 |
| Fli1 | 1.068989543 | Irs4 | -1.672753656 |
| Endog | 1.068893916 | Atp13a5 | -1.67188119 |
| Schip1 | 1.067163912 | Cdx2 | -1.671877604 |

| | | | |
|---|---|---|---|
| Slc38a3 | 1.066992652 | Serpinb1c | -1.671876502 |
| Naf1 | 1.062683542 | Mrap | -1.671875581 |
| Vstm4 | 1.062306562 | Gm364 | -1.671873195 |
| Timm8a1 | 1.062208494 | Abca17 | -1.671873103 |
| Syt1 | 1.061688023 | Lbp | -1.668751359 |
| Dnajc22 | 1.059891052 | Tmtc2 | -1.66733181 |
| Tomm20l | 1.058452875 | Myo1e | -1.667106773 |
| Sdc4 | 1.058184985 | Pml | -1.666256496 |
| Foxr2 | 1.057766256 | Csf3 | -1.666064638 |
| Cited4 | 1.056382318 | Tulp1 | -1.666064441 |
| Ppil6 | 1.056262246 | Abca12 | -1.666063758 |
| Tlr9 | 1.056160294 | Cd55 | -1.665827399 |
| Car2 | 1.052193254 | Cmbl | -1.664655709 |
| Cd101 | 1.052183839 | Idi2 | -1.664002011 |
| Fth1 | 1.050185308 | AF529169 | -1.662788765 |
| Sh2d7 | 1.049725432 | Tmem235 | -1.662196929 |
| Cox18 | 1.049510739 | Map3k7cl | -1.661130501 |
| Lrrc73 | 1.048915743 | Aox4 | -1.659776706 |
| Crhbp | 1.04814641 | Sstr4 | -1.659589528 |
| Sema4a | 1.046883324 | Angptl7 | -1.659555886 |
| Etl4 | 1.046756317 | Sox10 | -1.659043725 |
| Ube2l6 | 1.045333596 | Aim2 | -1.658959564 |
| Hs3st2 | 1.045300939 | Hspg2 | -1.658640546 |
| Rasd2 | 1.044977365 | Ttc12 | -1.658613591 |
| Cfh | 1.043077929 | Lama4 | -1.655919482 |
| Crot | 1.042344501 | Isg20 | -1.654181459 |
| Foxp1 | 1.041665479 | Olfml2a | -1.652504729 |
| Bcar3 | 1.041452162 | Il21 | -1.652453647 |
| Fam227b | 1.040455756 | Spata19 | -1.652453431 |
| Ccdc155 | 1.040363085 | Cnga2 | -1.652448552 |
| St3gal6 | 1.038215878 | 1810062G17Rik | -1.652448051 |
| Lgals9 | 1.03623318 | Spata20 | -1.652447605 |
| Sft2d1 | 1.03616714 | 4833420G17Rik | -1.651197086 |
| Gipr | 1.033542047 | Nrap | -1.650492666 |
| Dmrta2 | 1.033491289 | Spats1 | -1.649588715 |
| Ppp2r5a | 1.032399571 | Ciita | -1.645437535 |
| Lcp1 | 1.032163055 | Klf1 | -1.644987545 |
| Ptpn6 | 1.03172251 | Epyc | -1.643756085 |
| Cx3cr1 | 1.031678771 | Hist1h2ab | -1.643042588 |
| Mypn | 1.03030879 | Itga5 | -1.641403048 |
| Tulp2 | 1.028822477 | Lrrc8b | -1.637939927 |
| Rspo3 | 1.028720854 | Nhs | -1.635692932 |
| St6galnac6 | 1.028578582 | Wisp3 | -1.635178288 |
| Slc25a48 | 1.027806526 | Hist1h4m | -1.634596546 |
| Hhipl2 | 1.027097608 | Kif2c | -1.633734193 |
| Atp2a3 | 1.026051984 | AU022751 | -1.632036949 |
| Lemd1 | 1.024741085 | Sema3e | -1.631741924 |
| Gm14403 | 1.024504232 | Shisa6 | -1.630143171 |
| Htra2 | 1.02371048 | Kntc1 | -1.62929213 |
| Ehf | 1.021601342 | Ptpn22 | -1.627494937 |
| Lrrc10b | 1.021460622 | Vgll2 | -1.627075798 |
| Bdnf | 1.018742053 | Best2 | -1.626543332 |
| Kctd16 | 1.018203033 | Nusap1 | -1.622724128 |
| Panx2 | 1.017843446 | Fmn1 | -1.622501782 |
| Pld2 | 1.016597147 | Tmem108 | -1.620815019 |
| Frmd7 | 1.015926341 | Arid5b | -1.620564059 |
| Spry4 | 1.014436543 | Cyp2r1 | -1.61735635 |
| Sptb | 1.012915249 | Prrg3 | -1.617037669 |

| | | | |
|---|---|---|---|
| Gm9958 | 1.012573699 | Flrt1 | -1.614976233 |
| Trib1 | 1.010630274 | Emp2 | -1.614206255 |
| 4933425L06Rik | 1.00955282 | Matn2 | -1.6130492 |
| Osgin1 | 1.008467913 | Ppp1r17 | -1.612846884 |
| Cyp4f18 | 1.007104041 | Slc22a21 | -1.611549905 |
| Ndst3 | 1.006125233 | Spata21 | -1.611161578 |
| Kcnt1 | 1.005847806 | Galm | -1.610950944 |
| Ndufaf4 | 1.005274984 | Strc | -1.610288051 |
| Hcst | 1.004117867 | Hoxa3 | -1.608897537 |
| Cntnap3 | 1.004114891 | Plce1 | -1.606783035 |
| Cdh26 | 1.002839476 | Arhgef15 | -1.60466283 |
| Slc6a9 | 1.001698714 | Mc5r | -1.604257827 |
| Gpr82 | 1.001662637 | Fam71d | -1.604144936 |
| Hrh3 | 1.00104476 | Pcdh9 | -1.603650461 |
| | | Phldb3 | -1.602738157 |
| | | Cyp1b1 | -1.601491091 |
| | | Spon2 | -1.600802393 |
| | | Gpr182 | -1.600404981 |
| | | Syt13 | -1.600175391 |
| | | Pctp | -1.599749725 |
| | | Hmgb2 | -1.598784039 |
| | | 6330409D20Rik | -1.597963337 |
| | | Fmod | -1.5973336 |
| | | Stk33 | -1.596029421 |
| | | Slc22a3 | -1.59590918 |
| | | Pi4k2b | -1.59529508 |
| | | Incenp | -1.591984072 |
| | | Ust | -1.590815038 |
| | | Pdzrn3 | -1.589170625 |
| | | Srpx | -1.58864902 |
| | | St6gal2 | -1.588371011 |
| | | Ereg | -1.587948168 |
| | | Entpd8 | -1.585679235 |
| | | BC028528 | -1.585674849 |
| | | Bglap | -1.585306361 |
| | | Slc24a1 | -1.585164437 |
| | | Unc5c | -1.584479561 |
| | | Gen1 | -1.582994344 |
| | | Mfsd7a | -1.582566542 |
| | | Vmn2r1 | -1.581987826 |
| | | Ect2 | -1.581589734 |
| | | Cenpl | -1.576205923 |
| | | Cd109 | -1.575159605 |
| | | Tppp3 | -1.574591057 |
| | | Bub1 | -1.574586614 |
| | | Pabpc1l | -1.574197411 |
| | | Cdh6 | -1.573965175 |
| | | Tmem255a | -1.572451869 |
| | | Dnajb13 | -1.572206814 |
| | | Adamts1 | -1.57156028 |
| | | Prr15 | -1.571156382 |
| | | Jakmip3 | -1.570679262 |
| | | Map3k8 | -1.56771999 |
| | | Cartpt | -1.566680875 |
| | | Scara5 | -1.566222961 |
| | | Cabp5 | -1.563442259 |
| | | Bub1b | -1.562630478 |
| | | Vmn1r90 | -1.56231073 |

| | | | |
|---|---|---|---|
| | | Bmp4 | -1.56088832 |
| | | Zfp750 | -1.560311552 |
| | | Htr3a | -1.558001327 |
| | | Zfp811 | -1.557451534 |
| | | Pkd2 | -1.557247821 |
| | | Hmgb4 | -1.557158351 |
| | | Serpinb8 | -1.556913716 |
| | | Gucy1b2 | -1.556905416 |
| | | Pcdh20 | -1.555909617 |
| | | Hrct1 | -1.555568714 |
| | | Hrc | -1.553356496 |
| | | En1 | -1.553340247 |
| | | Adcy10 | -1.548391992 |
| | | Scube2 | -1.547119607 |
| | | Pram1 | -1.545422946 |
| | | Zfand4 | -1.54389002 |
| | | Chrnb4 | -1.541886511 |
| | | Stab2 | -1.541417694 |
| | | Arhgap19 | -1.540406894 |
| | | Vgll3 | -1.540107852 |
| | | Dcc | -1.539867674 |
| | | Tmem45a | -1.539139533 |
| | | Crct1 | -1.538327987 |
| | | Rnf152 | -1.537263392 |
| | | C4b | -1.537069034 |
| | | Pcdh17 | -1.536717794 |
| | | Aldob | -1.535953626 |
| | | Sgcd | -1.535223419 |
| | | Ube2c | -1.532230228 |
| | | Erich2 | -1.531178871 |
| | | Elf4 | -1.530192099 |
| | | Col4a1 | -1.528289451 |
| | | Mme | -1.526515603 |
| | | Tmem53 | -1.525350044 |
| | | Hist1h2ag | -1.52514875 |
| | | Mmp28 | -1.523910168 |
| | | Acot12 | -1.523631046 |
| | | Col1a2 | -1.523375204 |
| | | Tmem239 | -1.523034163 |
| | | Col6a4 | -1.523007372 |
| | | Ccnf | -1.522679776 |
| | | Kif20b | -1.522279988 |
| | | Nkx6-2 | -1.520870658 |
| | | Slco2a1 | -1.519445741 |
| | | Ppp1r32 | -1.518723797 |
| | | Col4a5 | -1.517677739 |
| | | Espl1 | -1.514229009 |
| | | Adamtsl3 | -1.513307936 |
| | | Tmeff2 | -1.512817147 |
| | | Aurka | -1.510577319 |
| | | Cd160 | -1.50720988 |
| | | Anln | -1.506531038 |
| | | Pcsk6 | -1.505024616 |
| | | Rnase4 | -1.504409343 |
| | | Gpr17 | -1.503603634 |
| | | Miip | -1.503313019 |
| | | Top2a | -1.501805458 |
| | | Rin1 | -1.500752114 |

| | | | |
|---|---|---|---|
| | | Sycp2 | -1.499866997 |
| | | Lrrc8c | -1.499530501 |
| | | Uba1y | -1.499032879 |
| | | Gm1141 | -1.49898099 |
| | | Fitm1 | -1.495673492 |
| | | Fancd2 | -1.49494176 |
| | | Sytl1 | -1.493950887 |
| | | Fam198b | -1.493905572 |
| | | Edn1 | -1.493123588 |
| | | Fbxo5 | -1.487879443 |
| | | Susd1 | -1.487610115 |
| | | Kcnt2 | -1.487346345 |
| | | Adam34 | -1.487286785 |
| | | Dupd1 | -1.487279766 |
| | | Slc46a2 | -1.487279444 |
| | | Ern2 | -1.487279256 |
| | | Hus1b | -1.487277932 |
| | | Cuzd1 | -1.487277523 |
| | | Slc5a4b | -1.487277226 |
| | | Apon | -1.487176349 |
| | | Pde11a | -1.487158271 |
| | | Sema4d | -1.486928624 |
| | | Tmprss6 | -1.486178282 |
| | | AI182371 | -1.486143756 |
| | | Ndc80 | -1.485653108 |
| | | Emp3 | -1.481574103 |
| | | Kdr | -1.481520547 |
| | | Ccdc36 | -1.481458056 |
| | | Epha2 | -1.478024584 |
| | | Cd27 | -1.477849647 |
| | | D6Ertd527e | -1.477849212 |
| | | Tep1 | -1.477176788 |
| | | Col3a1 | -1.477095867 |
| | | Melk | -1.476873265 |
| | | Mb | -1.474690208 |
| | | Dio2 | -1.47410394 |
| | | 1700123K08Rik | -1.473708723 |
| | | Dmbx1 | -1.473705699 |
| | | Depdc1b | -1.473030395 |
| | | Serpinf2 | -1.470477722 |
| | | Bcam | -1.470183963 |
| | | Dnaic2 | -1.469258044 |
| | | Armcx4 | -1.469111913 |
| | | Ago3 | -1.468968421 |
| | | Rtkn2 | -1.468050535 |
| | | Pcdhb2 | -1.467767303 |
| | | Zan | -1.467603199 |
| | | Parpbp | -1.467422086 |
| | | Hmmr | -1.466930534 |
| | | 4930452B06Rik | -1.466913778 |
| | | Sdr42e1 | -1.466305074 |
| | | Zdhhc23 | -1.466284266 |
| | | Pdlim1 | -1.466166695 |
| | | Fry | -1.466157816 |
| | | Plk1 | -1.46476716 |
| | | Tpx2 | -1.463786047 |
| | | Fam46b | -1.462147259 |
| | | Aspa | -1.462003965 |

| | | | |
|---|---|---|---|
| | | Rassf9 | -1.462003547 |
| | | Lrrc17 | -1.462002107 |
| | | Hsd3b6 | -1.462001174 |
| | | Avpr1b | -1.462001098 |
| | | Ush1c | -1.462000689 |
| | | Vmn2r57 | -1.46200065 |
| | | Tgm6 | -1.462000637 |
| | | Wfdc12 | -1.462000401 |
| | | Ropn1 | -1.461999679 |
| | | Fam129a | -1.459861104 |
| | | Anxa11 | -1.459553005 |
| | | D430041D05Rik | -1.459378315 |
| | | Ephb1 | -1.455719418 |
| | | Lgi3 | -1.455021901 |
| | | Edil3 | -1.454355926 |
| | | Epha4 | -1.454200479 |
| | | Krt24 | -1.453475722 |
| | | Fgl2 | -1.45298698 |
| | | Mmrn2 | -1.452907728 |
| | | Htr2c | -1.452580521 |
| | | Calhm2 | -1.451651368 |
| | | 4932443I19Rik | -1.450370451 |
| | | Nkx2-1 | -1.450311061 |
| | | Il17ra | -1.449958152 |
| | | AI467606 | -1.449196217 |
| | | Pcdh12 | -1.449195508 |
| | | Trim43b | -1.449194024 |
| | | Adam21 | -1.449170649 |
| | | Sdk1 | -1.44901769 |
| | | Hmgcll1 | -1.449017044 |
| | | Edar | -1.448723643 |
| | | S100a6 | -1.447653125 |
| | | Hs3st6 | -1.446812223 |
| | | Adamts14 | -1.446167045 |
| | | Thsd1 | -1.445067568 |
| | | Ankmy1 | -1.444361048 |
| | | Olfml2b | -1.443482741 |
| | | Kcnq3 | -1.443241452 |
| | | Muc15 | -1.443012104 |
| | | Kcnh7 | -1.442771144 |
| | | Kifc1 | -1.442654516 |
| | | Lmod1 | -1.440834753 |
| | | Agbl4 | -1.439384613 |
| | | Utrn | -1.439286641 |
| | | Slc12a2 | -1.438553006 |
| | | Ncapd2 | -1.438275569 |
| | | Tmsb15b2 | -1.43813951 |
| | | Esrrg | -1.437862289 |
| | | Sfn | -1.436990942 |
| | | Gm5475 | -1.436274731 |
| | | Wdr86 | -1.436274304 |
| | | I830077J02Rik | -1.436273703 |
| | | Tldc2 | -1.436272597 |
| | | Pla2g2e | -1.436271993 |
| | | Cdh15 | -1.436270881 |
| | | Card11 | -1.436269321 |
| | | Nox4 | -1.434923188 |
| | | Lrrk2 | -1.432776032 |

| | | | |
|---|---|---|---|
| | | Kcnk7 | -1.431592877 |
| | | Adam19 | -1.430305451 |
| | | Sec16b | -1.429612043 |
| | | Cep55 | -1.428305663 |
| | | Asic3 | -1.427114312 |
| | | Isoc2b | -1.427097537 |
| | | Golgb1 | -1.42493677 |
| | | Esrp2 | -1.424193932 |
| | | Qrich2 | -1.424149199 |
| | | Ildr1 | -1.423840887 |
| | | Pde8a | -1.423811501 |
| | | Ankrd28 | -1.421835288 |
| | | Gnat2 | -1.421237137 |
| | | Slc17a9 | -1.420711043 |
| | | Tnfrsf23 | -1.419870067 |
| | | Txnip | -1.419704102 |
| | | Prnd | -1.419346916 |
| | | Vasn | -1.418703111 |
| | | Myo7a | -1.416706733 |
| | | Nfe2 | -1.416532827 |
| | | Nipal3 | -1.416100857 |
| | | Cd209c | -1.415182879 |
| | | Ido2 | -1.414315062 |
| | | Grk1 | -1.4114333 |
| | | Dcdc2a | -1.411245081 |
| | | Ccdc171 | -1.410915838 |
| | | Zp1 | -1.410082276 |
| | | Pdyn | -1.410080174 |
| | | Nmu | -1.410078814 |
| | | Zscan10 | -1.410078424 |
| | | Slc4a1 | -1.41007838 |
| | | Psg29 | -1.410077725 |
| | | Ly6a | -1.410077431 |
| | | Zfp704 | -1.409783887 |
| | | Fbln1 | -1.406776992 |
| | | Id1 | -1.405755911 |
| | | Pon3 | -1.402596101 |
| | | Raet1d | -1.402370415 |
| | | Ptgir | -1.401101287 |
| | | Greb1 | -1.400573803 |
| | | Spag5 | -1.399539045 |
| | | Mcmdc2 | -1.398505129 |
| | | Lcor | -1.393455126 |
| | | Pak3 | -1.393394542 |
| | | Zfp933 | -1.3927209 |
| | | Nuak2 | -1.391735429 |
| | | Atp6v0a4 | -1.391351895 |
| | | Calr4 | -1.391252783 |
| | | Psg23 | -1.391086248 |
| | | Prr11 | -1.390499497 |
| | | Sulf2 | -1.3902749 |
| | | Nek2 | -1.390062042 |
| | | Cobll1 | -1.389571238 |
| | | Shroom1 | -1.389539161 |
| | | Birc5 | -1.389237349 |
| | | Nuggc | -1.387896776 |
| | | Cntn4 | -1.387448127 |
| | | Gpr179 | -1.387390833 |

| | | | |
|---|---|---|---|
| | | Ccdc110 | -1.387081064 |
| | | 4930486L24Rik | -1.386465074 |
| | | Zfp536 | -1.386012509 |
| | | D630045J12Rik | -1.383017198 |
| | | Guca1b | -1.382927393 |
| | | Slc1a1 | -1.382387352 |
| | | Pdcd1lg2 | -1.381617658 |
| | | Gpr161 | -1.380019612 |
| | | Tmsb15l | -1.378543283 |
| | | Tert | -1.377647357 |
| | | Zpld1 | -1.377634128 |
| | | Crygs | -1.376849684 |
| | | Akap6 | -1.375253314 |
| | | Tes | -1.375011778 |
| | | Heg1 | -1.374263036 |
| | | Klf8 | -1.374243075 |
| | | Slc16a8 | -1.373093615 |
| | | Serinc2 | -1.373001859 |
| | | Smoc2 | -1.372689292 |
| | | Acta2 | -1.372288629 |
| | | Slit1 | -1.371993464 |
| | | Mfsd9 | -1.371737286 |
| | | Olfr613 | -1.370486851 |
| | | Lbx2 | -1.369265849 |
| | | Herc6 | -1.367940275 |
| | | Lhfpl2 | -1.36791999 |
| | | Fat1 | -1.365776705 |
| | | Rtn4rl2 | -1.364267132 |
| | | Tmem178b | -1.363737566 |
| | | Ryr1 | -1.36371717 |
| | | Cpa4 | -1.36296864 |
| | | Hspb1 | -1.362539612 |
| | | Tgfbr3 | -1.362293235 |
| | | Slc25a24 | -1.361752643 |
| | | Zfp941 | -1.361578035 |
| | | Notch1 | -1.361248267 |
| | | Knstrn | -1.359291044 |
| | | Agbl2 | -1.358873841 |
| | | Npffr1 | -1.357403439 |
| | | Eml2 | -1.357378903 |
| | | Tmc1 | -1.355393088 |
| | | Kif18b | -1.354749483 |
| | | Cbln2 | -1.352398874 |
| | | Bmf | -1.352019587 |
| | | Syt14 | -1.3512878 |
| | | Selp | -1.34981324 |
| | | Slc19a3 | -1.349812837 |
| | | C1ql3 | -1.34939858 |
| | | Best1 | -1.347862967 |
| | | Astn2 | -1.347064083 |
| | | Fzd5 | -1.346871338 |
| | | Polq | -1.345465757 |
| | | Pde1a | -1.344827567 |
| | | Psmd9 | -1.344820176 |
| | | Syne1 | -1.344262974 |
| | | Macf1 | -1.343614383 |
| | | Sema3d | -1.343467434 |
| | | Fam228a | -1.343060772 |

| | | | |
|---|---|---|---|
| | | Uba7 | -1.34293325 |
| | | Zfp773 | -1.342499964 |
| | | Prox1 | -1.342359134 |
| | | Brdt | -1.340730548 |
| | | Acvrl1 | -1.339487757 |
| | | Rps6ka1 | -1.339235446 |
| | | Ephb2 | -1.339064155 |
| | | 4933402D24Rik | -1.338616408 |
| | | Pcdhb21 | -1.338458919 |
| | | Hipk4 | -1.334686466 |
| | | Slc27a5 | -1.334151546 |
| | | Ebf1 | -1.333779474 |
| | | Hydin | -1.33154862 |
| | | Thsd7a | -1.330654772 |
| | | Catsperd | -1.330071175 |
| | | Ttk | -1.329940467 |
| | | Nkx2-2 | -1.32944973 |
| | | Cyp2j8 | -1.328832 |
| | | Ccdc18 | -1.328550985 |
| | | Socs3 | -1.328507391 |
| | | Plin4 | -1.328309819 |
| | | Lrg1 | -1.328156222 |
| | | Slc44a5 | -1.327724301 |
| | | Fv1 | -1.327577598 |
| | | Cdh19 | -1.327344031 |
| | | Rbm47 | -1.327232057 |
| | | Inhbc | -1.327168752 |
| | | Cdk5rap2 | -1.327029091 |
| | | H2-T24 | -1.326412529 |
| | | Pcdhb15 | -1.324718483 |
| | | Pdgfrb | -1.324135922 |
| | | Kif5a | -1.32392894 |
| | | Lrrc69 | -1.323853972 |
| | | Nrcam | -1.323680831 |
| | | 5730507C01Rik | -1.322994105 |
| | | Jag1 | -1.322970909 |
| | | Ttll13 | -1.322432799 |
| | | Esco2 | -1.31925522 |
| | | Tet2 | -1.318303092 |
| | | Akap9 | -1.317023776 |
| | | Fam46c | -1.31677475 |
| | | Ston2 | -1.314049278 |
| | | Trpc4 | -1.313898252 |
| | | Il1rapl2 | -1.31389066 |
| | | Ghrh | -1.313859673 |
| | | Cyfip2 | -1.313567858 |
| | | Eml1 | -1.313333574 |
| | | Slc38a6 | -1.312848307 |
| | | G2e3 | -1.311456449 |
| | | Mitf | -1.310747997 |
| | | Cdh11 | -1.309003654 |
| | | Dixdc1 | -1.307971189 |
| | | Itpr3 | -1.306521756 |
| | | Plaur | -1.306507506 |
| | | A730017C20Rik | -1.306222597 |
| | | Pnma3 | -1.305230862 |
| | | Ska3 | -1.304455187 |
| | | Lctl | -1.303887955 |

| | | | |
|---|---|---|---|
| | | Brip1 | -1.3037254 |
| | | Pot1b | -1.302860426 |
| | | Dok7 | -1.302799263 |
| | | Nrp2 | -1.301954408 |
| | | Prss8 | -1.301527407 |
| | | Rpgrip1l | -1.301302634 |
| | | Strn | -1.300557271 |
| | | Lgals1 | -1.298889344 |
| | | Syt2 | -1.297064574 |
| | | Cep89 | -1.29686319 |
| | | Cdca3 | -1.29652914 |
| | | Poc1a | -1.295471955 |
| | | Gm6377 | -1.294925776 |
| | | Defb20 | -1.294922269 |
| | | Cxcl17 | -1.294921783 |
| | | 4931409K22Rik | -1.294920113 |
| | | Prps1l1 | -1.294919626 |
| | | Rln3 | -1.294917912 |
| | | Ppp3r2 | -1.292348552 |
| | | Trim69 | -1.288883287 |
| | | Hhatl | -1.288298975 |
| | | Galns | -1.287730846 |
| | | Loxl4 | -1.287672625 |
| | | H2-Q4 | -1.286060486 |
| | | Jsrp1 | -1.286059204 |
| | | Cdkn3 | -1.285647975 |
| | | Nox1 | -1.28233825 |
| | | Fat2 | -1.282338047 |
| | | Mid1 | -1.28201619 |
| | | Erbb3 | -1.281349867 |
| | | Cep112 | -1.280773914 |
| | | Lrrn4cl | -1.280440068 |
| | | S100a2 | -1.279269036 |
| | | Dbf4 | -1.278016093 |
| | | Troap | -1.277864004 |
| | | Slc16a12 | -1.277859857 |
| | | Islr2 | -1.277756726 |
| | | Trpm3 | -1.277665551 |
| | | Ccdc33 | -1.276623183 |
| | | Qpct | -1.27633802 |
| | | Fndc9 | -1.272972633 |
| | | Cd1d1 | -1.272769327 |
| | | Cenpp | -1.269960365 |
| | | Mslnl | -1.269647612 |
| | | Atp4b | -1.269645983 |
| | | Tmem182 | -1.269645043 |
| | | Fasl | -1.269644804 |
| | | Olfr1420 | -1.269644736 |
| | | Ugt2b37 | -1.269644278 |
| | | Dsc1 | -1.269643988 |
| | | Olfr90 | -1.269643152 |
| | | Mrgprx2 | -1.269641946 |
| | | Kif22 | -1.268295991 |
| | | Mis18bp1 | -1.266655719 |
| | | Dnaaf3 | -1.263563144 |
| | | Cdc25c | -1.262450712 |
| | | Gjb3 | -1.261118922 |
| | | Pecam1 | -1.260615041 |

| | | | |
|---|---|---|---|
| | | Ltb | -1.259844661 |
| | | Sned1 | -1.259479844 |
| | | Slc10a1 | -1.258526143 |
| | | Ptprj | -1.257353805 |
| | | Ano7 | -1.256837361 |
| | | Papln | -1.256240993 |
| | | Kazn | -1.255426792 |
| | | Kcnq1 | -1.254146427 |
| | | Adam4 | -1.254020181 |
| | | Pth1r | -1.253404972 |
| | | Chrm2 | -1.2532135 |
| | | Vdr | -1.253082643 |
| | | Ncapg | -1.252975844 |
| | | Crb2 | -1.252862052 |
| | | Lrig3 | -1.25110054 |
| | | Suco | -1.248085298 |
| | | Acss3 | -1.247641309 |
| | | Cast | -1.24659427 |
| | | Zfp369 | -1.245595862 |
| | | Mast4 | -1.24556239 |
| | | Rsph4a | -1.245145711 |
| | | Slc23a1 | -1.245108028 |
| | | Ccdc82 | -1.244637962 |
| | | Anxa2 | -1.244203972 |
| | | Vill | -1.243923651 |
| | | Arhgap8 | -1.243917318 |
| | | Crip3 | -1.242976238 |
| | | Hdac1 | -1.24270228 |
| | | Tspan18 | -1.242426117 |
| | | Prc1 | -1.242162569 |
| | | 1700022I11Rik | -1.241242303 |
| | | Racgap1 | -1.241052825 |
| | | Mmp11 | -1.24094393 |
| | | Atp6v1c2 | -1.240667962 |
| | | Zfp618 | -1.240402285 |
| | | Gcnt4 | -1.240296406 |
| | | Sdk2 | -1.236330605 |
| | | Cpne1 | -1.235818607 |
| | | Lamc2 | -1.23572863 |
| | | Ccng2 | -1.235555583 |
| | | Decr2 | -1.235544146 |
| | | Sorbs2 | -1.234992411 |
| | | Syn3 | -1.233518748 |
| | | Llgl2 | -1.233264185 |
| | | Pcdh19 | -1.232871573 |
| | | Rasgrp3 | -1.232450216 |
| | | Slc26a5 | -1.231430852 |
| | | Cxcr6 | -1.231426781 |
| | | Sh3d19 | -1.231305954 |
| | | Iqgap1 | -1.231255756 |
| | | Best3 | -1.230880502 |
| | | Prrg4 | -1.230588586 |
| | | Hsf2bp | -1.227789894 |
| | | Abcb1a | -1.227789453 |
| | | Ckap5 | -1.226926412 |
| | | Kcnj13 | -1.226307687 |
| | | Stmn2 | -1.225962765 |
| | | Gsta3 | -1.224821909 |

| | | | |
|---|---|---|---|
| | | Hoxb5 | -1.224246614 |
| | | Spdl1 | -1.224074387 |
| | | Cchcr1 | -1.223802177 |
| | | Col11a1 | -1.222999091 |
| | | Opn3 | -1.222870845 |
| | | Sec31b | -1.221877027 |
| | | Mylpf | -1.221661975 |
| | | Snapc1 | -1.221506712 |
| | | Rad51b | -1.221433216 |
| | | Baz2b | -1.221053821 |
| | | Itgbl1 | -1.220134529 |
| | | Flnb | -1.219868018 |
| | | Fam229a | -1.217723846 |
| | | Snx31 | -1.217722457 |
| | | Serpini2 | -1.217722224 |
| | | Sult5a1 | -1.21772194 |
| | | Hsd17b14 | -1.217720937 |
| | | Kctd14 | -1.216205144 |
| | | Mnd1 | -1.215818865 |
| | | Nxph3 | -1.215267952 |
| | | Tcf7l2 | -1.214737264 |
| | | Emb | -1.214429958 |
| | | Grhl2 | -1.214187865 |
| | | Dusp23 | -1.213954344 |
| | | Kcnmb1 | -1.212258501 |
| | | Pcnt | -1.212032387 |
| | | Cdk19 | -1.211214083 |
| | | Nek3 | -1.210086956 |
| | | Unc13b | -1.208590087 |
| | | Aurkb | -1.207345303 |
| | | Kidins220 | -1.207215012 |
| | | Mia3 | -1.207155372 |
| | | Zkscan7 | -1.206287714 |
| | | Lrrtm2 | -1.20426901 |
| | | Odf2 | -1.203433106 |
| | | Rictor | -1.203092313 |
| | | 1700007K09Rik | -1.202580224 |
| | | Rgs16 | -1.202576673 |
| | | Xkr7 | -1.20252884 |
| | | Grik1 | -1.202225388 |
| | | Evi2a | -1.20188828 |
| | | Slc12a7 | -1.200488772 |
| | | Col11a2 | -1.19983102 |
| | | Bnc2 | -1.199341157 |
| | | Spata13 | -1.197717881 |
| | | Manba | -1.194913729 |
| | | Tmod2 | -1.19488446 |
| | | Wfikkn2 | -1.194023061 |
| | | Bmp15 | -1.193859019 |
| | | Epn3 | -1.193856661 |
| | | Tbx21 | -1.193856022 |
| | | Cntnap5c | -1.193854192 |
| | | Omp | -1.193810832 |
| | | Aph1c | -1.193343708 |
| | | Rnf122 | -1.193319577 |
| | | Disp2 | -1.193202295 |
| | | Tmem63c | -1.193081688 |
| | | Epha6 | -1.192731626 |

| | | | |
|---|---|---|---|
| | | Npm2 | -1.192345953 |
| | | Alas2 | -1.192202533 |
| | | Tfcp2l1 | -1.19204589 |
| | | Tenm2 | -1.192016706 |
| | | Scube3 | -1.191463138 |
| | | Pik3r3 | -1.191104517 |
| | | Nlgn1 | -1.191027685 |
| | | Acyp1 | -1.18964793 |
| | | Slc43a3 | -1.187349213 |
| | | 1700020D05Rik | -1.186941805 |
| | | Camk4 | -1.185497043 |
| | | Rab9b | -1.184108991 |
| | | Robo2 | -1.183895002 |
| | | Rnf125 | -1.183840187 |
| | | E2f7 | -1.183704364 |
| | | Icosl | -1.183630601 |
| | | Pcdhb3 | -1.182626493 |
| | | Creb3l4 | -1.182587338 |
| | | Hcn1 | -1.181516013 |
| | | Hist1h4c | -1.181010615 |
| | | Frmd5 | -1.180731087 |
| | | Slc6a3 | -1.179888359 |
| | | Plin5 | -1.17910552 |
| | | Ecm1 | -1.178823005 |
| | | Plag1 | -1.178423514 |
| | | Dusp16 | -1.178298095 |
| | | Ccna2 | -1.178174141 |
| | | Zeb2 | -1.177789452 |
| | | Flrt2 | -1.177194514 |
| | | Cmah | -1.176885402 |
| | | Hcrtr2 | -1.176881659 |
| | | Cdh18 | -1.176811018 |
| | | Helb | -1.175078191 |
| | | Tmtc4 | -1.173186711 |
| | | Henmt1 | -1.172636222 |
| | | Fbln7 | -1.169095286 |
| | | Ikbke | -1.168999678 |
| | | Runx1 | -1.168422246 |
| | | Ckap2l | -1.168245784 |
| | | Agtrap | -1.168161998 |
| | | Bhlha15 | -1.167383216 |
| | | Cobl | -1.167243539 |
| | | Piezo1 | -1.167241841 |
| | | Zranb3 | -1.167142072 |
| | | Gabrg3 | -1.166458032 |
| | | Xrra1 | -1.166272355 |
| | | Tnfaip2 | -1.166069754 |
| | | Slc38a4 | -1.165664266 |
| | | Cep350 | -1.165041879 |
| | | Otud7b | -1.164590729 |
| | | Vat1l | -1.163972233 |
| | | Stil | -1.162903063 |
| | | Sorcs1 | -1.162867337 |
| | | Oas3 | -1.162822984 |
| | | Pla2g2f | -1.162819464 |
| | | Mfsd6l | -1.162818506 |
| | | Zfp819 | -1.162818213 |
| | | A530016L24Rik | -1.16253565 |

| | | | |
|---|---|---|---|
| | | Hspb11 | -1.161174404 |
| | | Rffl | -1.161085037 |
| | | Efcab11 | -1.160927049 |
| | | Grin3a | -1.160205877 |
| | | Lgals3 | -1.158955181 |
| | | Trim30d | -1.15813147 |
| | | Armc2 | -1.157860247 |
| | | Gm14434 | -1.157399984 |
| | | Gm4724 | -1.157399984 |
| | | Pcdhb16 | -1.156910851 |
| | | Shroom3 | -1.156574659 |
| | | Gal3st1 | -1.156545927 |
| | | Efnb2 | -1.156391978 |
| | | D130040H23Rik | -1.15632073 |
| | | Tnnt1 | -1.155661794 |
| | | Tmc3 | -1.154751387 |
| | | Prox2 | -1.15452822 |
| | | Anxa3 | -1.154203184 |
| | | Fbn1 | -1.154146531 |
| | | Cep72 | -1.153456903 |
| | | Plxna2 | -1.153432714 |
| | | Pdgfra | -1.152686803 |
| | | Kcns3 | -1.151779444 |
| | | Pcdh7 | -1.151746128 |
| | | Amhr2 | -1.151439183 |
| | | Chrnb1 | -1.150709324 |
| | | Fam161a | -1.150689456 |
| | | Poln | -1.149224464 |
| | | Atp2a1 | -1.149124475 |
| | | Snai1 | -1.148968517 |
| | | Sfi1 | -1.14719006 |
| | | Cdkl2 | -1.146873011 |
| | | Tgfb1i1 | -1.146086376 |
| | | Cdca8 | -1.146081334 |
| | | Cdr1 | -1.1459257 |
| | | Zfp433 | -1.145646511 |
| | | Fga | -1.14536821 |
| | | Myo1a | -1.144773693 |
| | | Cnr1 | -1.144341227 |
| | | Catsperg1 | -1.143309707 |
| | | Ttbk2 | -1.143198918 |
| | | Npy | -1.143105872 |
| | | Gm6525 | -1.141938203 |
| | | Slc16a7 | -1.141548759 |
| | | Shcbp1 | -1.140666721 |
| | | Ephb6 | -1.140481551 |
| | | Rnf144b | -1.139883584 |
| | | Sez6l | -1.139808687 |
| | | Ccbe1 | -1.139795665 |
| | | Cabyr | -1.139167594 |
| | | Mysm1 | -1.137977389 |
| | | 9130008F23Rik | -1.137816108 |
| | | Myh11 | -1.137754251 |
| | | Xrn1 | -1.136968707 |
| | | Sln | -1.136810549 |
| | | Hist1h3c | -1.136348303 |
| | | Sema4c | -1.13627214 |
| | | Cpxm2 | -1.135740078 |

| | | | |
|---|---|---|---|
| | | Ccdc8 | -1.135240824 |
| | | Ccin | -1.134811354 |
| | | Rttn | -1.133468934 |
| | | Htr1b | -1.133431106 |
| | | Eif4ebp3 | -1.13321853 |
| | | Trp63 | -1.1319584 |
| | | 1700034I23Rik | -1.131749464 |
| | | Sv2b | -1.131624507 |
| | | Slc29a3 | -1.131467744 |
| | | Rhbdf2 | -1.131260174 |
| | | Zfp772 | -1.131213943 |
| | | Cercam | -1.129841215 |
| | | Pnma2 | -1.128677086 |
| | | Plp2 | -1.128662374 |
| | | Il6ra | -1.128462597 |
| | | Stap2 | -1.128097857 |
| | | Pou5f2 | -1.128014882 |
| | | Artn | -1.127448846 |
| | | Fam83d | -1.127167721 |
| | | Capn1 | -1.126852919 |
| | | Pak1 | -1.126844306 |
| | | Nckap5 | -1.126523837 |
| | | Rab11fip1 | -1.126397652 |
| | | Micall2 | -1.126006937 |
| | | Cldn1 | -1.124339005 |
| | | Proca1 | -1.124220163 |
| | | Trim30a | -1.123345601 |
| | | Ankrd34b | -1.123315721 |
| | | Ptprb | -1.12134348 |
| | | Gria4 | -1.120871355 |
| | | Plekhg3 | -1.12070862 |
| | | Zbtb37 | -1.120230937 |
| | | Slc8a2 | -1.119733089 |
| | | Sesn3 | -1.119388337 |
| | | Ly6g6f | -1.118988581 |
| | | Spc25 | -1.118945713 |
| | | Cep192 | -1.117273337 |
| | | Ccdc148 | -1.117216956 |
| | | Tnnc2 | -1.116920038 |
| | | Bach2 | -1.116394699 |
| | | Ada | -1.11624422 |
| | | Col1a1 | -1.115763552 |
| | | Atp10a | -1.115548046 |
| | | Gdf5 | -1.11534853 |
| | | Fam134b | -1.115314 |
| | | Dpysl4 | -1.114846865 |
| | | Prdm10 | -1.114553328 |
| | | Fam46a | -1.114028144 |
| | | Igf2r | -1.113982416 |
| | | Myom1 | -1.113910954 |
| | | Olfr1349 | -1.1136593 |
| | | Nid2 | -1.113396333 |
| | | Ero1lb | -1.1129455 |
| | | Ninl | -1.111843229 |
| | | Dok6 | -1.110927539 |
| | | Gpha2 | -1.109747854 |
| | | Col4a3 | -1.107429085 |
| | | Gng13 | -1.107247716 |

| | | | |
|---|---|---|---|
| | | Usp40 | -1.107174351 |
| | | Iqcc | -1.106825389 |
| | | Dnm3 | -1.106700844 |
| | | 4932438A13Rik | -1.106454646 |
| | | Slc37a1 | -1.105951153 |
| | | Tapbpl | -1.104387149 |
| | | Rapgef5 | -1.103958154 |
| | | Cep250 | -1.103785086 |
| | | Psd2 | -1.103581131 |
| | | Capsl | -1.103348318 |
| | | Cyp26c1 | -1.103070231 |
| | | Cadm1 | -1.102808809 |
| | | Ifnz | -1.102131574 |
| | | Clcn5 | -1.101537253 |
| | | Zfp804b | -1.101529821 |
| | | Actn1 | -1.100832289 |
| | | Galnt3 | -1.100761923 |
| | | Gxylt1 | -1.100715399 |
| | | Stx2 | -1.100594708 |
| | | Cacnb2 | -1.10025261 |
| | | Gm5415 | -1.100238291 |
| | | Prss35 | -1.100171078 |
| | | Zfp791 | -1.099955181 |
| | | 1700093K21Rik | -1.099586138 |
| | | Ang | -1.099367886 |
| | | Apobec2 | -1.099258911 |
| | | Anpep | -1.098124354 |
| | | Myo6 | -1.097827215 |
| | | Nradd | -1.097713146 |
| | | Neurl1b | -1.097212087 |
| | | Trip11 | -1.097181817 |
| | | Ap1s3 | -1.096781075 |
| | | Adamts18 | -1.096529872 |
| | | Kcnj9 | -1.09638918 |
| | | Tmem2 | -1.095287958 |
| | | Naip1 | -1.094478686 |
| | | Lrrc72 | -1.094476762 |
| | | Spaca5 | -1.094475465 |
| | | H2-DMb1 | -1.094474925 |
| | | Glra4 | -1.094473637 |
| | | Cpeb4 | -1.092711103 |
| | | Scml2 | -1.091833755 |
| | | Cep85l | -1.09112182 |
| | | Dach2 | -1.09080411 |
| | | Ankrd53 | -1.090509262 |
| | | St8sia3 | -1.090188759 |
| | | Lrig1 | -1.090074137 |
| | | Adamtsl1 | -1.089657443 |
| | | Rprm | -1.089463165 |
| | | Smtn | -1.089340425 |
| | | Ctrl | -1.089247853 |
| | | Tekt2 | -1.087419196 |
| | | Gcnt7 | -1.087383071 |
| | | Ska1 | -1.087229154 |
| | | 4930550C14Rik | -1.086912963 |
| | | Hpse | -1.086837092 |
| | | Gal | -1.086779781 |
| | | Cd247 | -1.086777494 |

| | | | |
|---|---|---|---|
| | | Rab33a | -1.08674746 |
| | | Tmem245 | -1.086387866 |
| | | Slc8a1 | -1.086126166 |
| | | Fam129b | -1.086095195 |
| | | Trim15 | -1.085679117 |
| | | Ubxn10 | -1.085512952 |
| | | Parm1 | -1.084889085 |
| | | Chst3 | -1.084694918 |
| | | Bora | -1.083561643 |
| | | Fam221b | -1.083142993 |
| | | Gsta1 | -1.081102942 |
| | | Mcidas | -1.081100415 |
| | | Hipk2 | -1.07948944 |
| | | Wfdc3 | -1.079041653 |
| | | Tulp4 | -1.078228912 |
| | | Cep152 | -1.078091302 |
| | | Crb3 | -1.077930165 |
| | | Pnp2 | -1.077235214 |
| | | Ncapg2 | -1.076981891 |
| | | Tab3 | -1.075554454 |
| | | Efr3b | -1.075408289 |
| | | Gas2l3 | -1.075234051 |
| | | Stard6 | -1.074992598 |
| | | Abcb4 | -1.074907879 |
| | | Atp8a1 | -1.07410527 |
| | | Gng8 | -1.073917145 |
| | | Tcf7l1 | -1.073671333 |
| | | Pkn3 | -1.073611451 |
| | | Foxf2 | -1.073207916 |
| | | Psen2 | -1.072794091 |
| | | Tspan15 | -1.072160127 |
| | | Plch1 | -1.071356968 |
| | | Pcdhb7 | -1.071313116 |
| | | Reep4 | -1.070892352 |
| | | Ggt5 | -1.069989063 |
| | | Olfr267 | -1.069987105 |
| | | Acr | -1.069956735 |
| | | Pls3 | -1.069934699 |
| | | Gsg1 | -1.068657938 |
| | | Phka2 | -1.06730491 |
| | | Adamts19 | -1.06707255 |
| | | Man1a | -1.065962872 |
| | | Nalcn | -1.063988225 |
| | | Nppb | -1.063435277 |
| | | Hebp2 | -1.063231578 |
| | | Tstd3 | -1.060972452 |
| | | Dusp4 | -1.060184297 |
| | | Frem1 | -1.060013685 |
| | | Gprin3 | -1.059981128 |
| | | Pdcd4 | -1.059337804 |
| | | Cnp | -1.059138293 |
| | | Podxl | -1.05911385 |
| | | Pdgfa | -1.058625578 |
| | | Lhx6 | -1.058197212 |
| | | Trpv2 | -1.057566402 |
| | | Lnpep | -1.056447173 |
| | | Map7 | -1.055982284 |
| | | Csmd3 | -1.055379263 |

| | | | |
|---|---|---|---|
| | | Hyls1 | -1.055099386 |
| | | Pate4 | -1.054897472 |
| | | Tecta | -1.054353557 |
| | | Grm1 | -1.05263304 |
| | | Trim56 | -1.051672386 |
| | | Npnt | -1.05076084 |
| | | Nsun6 | -1.049890338 |
| | | Scarb2 | -1.04967292 |
| | | Ccdc77 | -1.049653997 |
| | | Dnajc6 | -1.049565783 |
| | | Lcorl | -1.048507059 |
| | | Kif24 | -1.048453131 |
| | | Mmp25 | -1.047507185 |
| | | Fam160a1 | -1.047323842 |
| | | Clec1a | -1.046964588 |
| | | Mab21l2 | -1.046963675 |
| | | Mfsd6 | -1.0469196 |
| | | Igfbpl1 | -1.046552172 |
| | | Cd9 | -1.046404052 |
| | | Mdm1 | -1.045800085 |
| | | Unc5d | -1.045026353 |
| | | Secisbp2l | -1.044997186 |
| | | Atf6 | -1.044147445 |
| | | Kif21a | -1.044139411 |
| | | Amz1 | -1.043622847 |
| | | Obscn | -1.042843668 |
| | | Pros1 | -1.042794717 |
| | | Gli3 | -1.042337758 |
| | | Apold1 | -1.042296888 |
| | | Mef2b | -1.041798534 |
| | | Dync1h1 | -1.041533433 |
| | | Tacc3 | -1.041370878 |
| | | 5730480H06Rik | -1.041062461 |
| | | Celsr1 | -1.041039074 |
| | | Pcdhb5 | -1.040898383 |
| | | Pear1 | -1.040881431 |
| | | Hist2h4 | -1.039772473 |
| | | Rdh12 | -1.039021311 |
| | | Dock6 | -1.038723488 |
| | | 4930503L19Rik | -1.038688233 |
| | | Ggta1 | -1.037621111 |
| | | Cda | -1.036995896 |
| | | Tnfaip8 | -1.035673858 |
| | | Gm7694 | -1.033741126 |
| | | Nphp3 | -1.032996109 |
| | | Lama3 | -1.032069296 |
| | | Rdh16 | -1.031888627 |
| | | Zbp1 | -1.031886612 |
| | | 4833427G06Rik | -1.031808544 |
| | | Kirrel3 | -1.031076855 |
| | | Nnmt | -1.030884394 |
| | | Wnt5b | -1.029892173 |
| | | H2-M10.2 | -1.029697368 |
| | | Fstl5 | -1.029455654 |
| | | Hc | -1.029030434 |
| | | Fstl4 | -1.028613214 |
| | | Gab3 | -1.028242377 |
| | | Lif | -1.028023528 |

| | | | |
|---|---|---|---|
| | | Slc44a3 | -1.027794319 |
| | | 9530053A07Rik | -1.026697803 |
| | | Fbxo24 | -1.026581592 |
| | | 9030624G23Rik | -1.02631674 |
| | | Rdh5 | -1.025875551 |
| | | Nudt12 | -1.025825029 |
| | | 1810041L15Rik | -1.025287612 |
| | | B3gnt7 | -1.025119209 |
| | | Cps1 | -1.02435007 |
| | | Cenpa | -1.023775422 |
| | | Sdcbp2 | -1.023526429 |
| | | Herc1 | -1.022652859 |
| | | Frem2 | -1.022393786 |
| | | Ccdc50 | -1.02235126 |
| | | Klhl28 | -1.022289415 |
| | | Plcxd3 | -1.022272588 |
| | | Eif2ak3 | -1.02208401 |
| | | Wisp1 | -1.022034526 |
| | | Slc27a3 | -1.021849345 |
| | | Hdx | -1.021698158 |
| | | Als2cl | -1.021507875 |
| | | Sun2 | -1.021313891 |
| | | Chml | -1.021003228 |
| | | Reck | -1.020812145 |
| | | Arrdc3 | -1.02071469 |
| | | Pklr | -1.019614142 |
| | | D430019H16Rik | -1.019576623 |
| | | Pcdh15 | -1.019521038 |
| | | Wnt2b | -1.01913201 |
| | | Ccnb2 | -1.019084597 |
| | | Grik3 | -1.01835419 |
| | | Klrg2 | -1.017224418 |
| | | Spatc1l | -1.016774831 |
| | | Iqcj | -1.016772675 |
| | | 1700019A02Rik | -1.016769496 |
| | | Atp7a | -1.01624918 |
| | | Eln | -1.016166486 |
| | | Lrig2 | -1.016066585 |
| | | Cpd | -1.01557842 |
| | | Palld | -1.01524609 |
| | | Ccdc146 | -1.01439719 |
| | | Slx4ip | -1.014322597 |
| | | Gabre | -1.014092863 |
| | | 1700029H14Rik | -1.014091332 |
| | | Col5a2 | -1.013795224 |
| | | Frrs1 | -1.012543837 |
| | | Tgfbi | -1.012242495 |
| | | Gabrb2 | -1.01217463 |
| | | Atl1 | -1.012099934 |
| | | Scyl2 | -1.01162434 |
| | | Mtus1 | -1.011372059 |
| | | Tekt1 | -1.011287623 |
| | | Duxbl1 | -1.01117077 |
| | | Duxbl3 | -1.011169759 |
| | | Duxbl2 | -1.011169759 |
| | | Rtn4 | -1.010422672 |
| | | 3300002I08Rik | -1.010264393 |
| | | Zbed6 | -1.009960212 |

| | | | |
|---|---|---|---|
| | | St8sia2 | -1.009788351 |
| | | Anxa7 | -1.00957039 |
| | | Tmbim1 | -1.009244727 |
| | | Tgfbr1 | -1.009142634 |
| | | Myh10 | -1.009078683 |
| | | CK137956 | -1.008866355 |
| | | Mansc1 | -1.008843471 |
| | | Kif18a | -1.008667804 |
| | | Lmbrd2 | -1.00844331 |
| | | Daam1 | -1.008373426 |
| | | Xkr5 | -1.008278653 |
| | | Acad12 | -1.007910595 |
| | | Mroh2a | -1.007829601 |
| | | Cdc20 | -1.007226401 |
| | | Grm5 | -1.006914693 |
| | | Pcdhb19 | -1.004841124 |
| | | Plau | -1.004265719 |
| | | Creb3l2 | -1.004136056 |
| | | Ell2 | -1.003829292 |
| | | Slc4a5 | -1.003825839 |
| | | Tbce | -1.003770904 |
| | | Atp11b | -1.003149541 |
| | | Htr4 | -1.002983016 |
| | | Cdc25b | -1.002612127 |
| | | Olfr70 | -1.001499296 |
| | | Utp14b | -1.001002777 |
| | | Kif20a | -1.000756757 |

**Supplementary Table 3. Primer sequences for qRT-PCR assay of target genes.**

| Gene | Forward primer | Reverse primer |
|---|---|---|
| *Ccna2* | TGGATGGCAGTTTTGAATCACC | CCCTAAGGTACGTGTGAATGTC |
| *Ccnb1* | GCCAAGAGCCATGTGACTATC | CAGAGCTGGTACTTTGGTGTTC |
| *Ccnf* | AGAGACTGAATACGGGTTCTGA | TCCCAAGCAGTGTAGTATGGAA |
| *Ccng2* | AGGGGTTCAGCTTTTCGGATT | AGTGTTATCATTCTCCGGGGTAG |
| *Cdc25b* | TCCGATCCTTACCAGTGAGG | GGGCAGAGCTGGAATGAGG |
| *Cdc25c* | GGCAAACCTAAGCATTCTGTCG | CCAGAGGTCCAGATGAATCCA |
| *Cdca3* | CTGAGCGAAGTATTGGAGACAG | CTGCGGATTGTTTGGCTTCC |
| *Cdk19* | GGTCAAGCCTGACAGCAAAGT | TTCCTGGAAGTAAGGGTCCTG |
| *Cldn11* | ATGGTAGCCACTTGCCTTCAG | AGTTCGTCCATTTTTCGGCAG |
| *Enpp2* | TTTGCACTATGCCAACAATCGG | GGAGGCACTTTAGTCCTGTACTT |
| *Kif11* | GGCTGGTATAATTCCACGCAC | CCGGGGATCATCAAACATCTG |
| *Mbp* | GCAGCCAGCACCACTCTTGA | CAGCCGAGGTCCCATTGTTC |
| *Myrf* | CCTGTGTCCGTGGTACTGTG | TCACACAGGCGGTAGAAGTG |
| *Cnp* | TTTACCCGCAAAAGCCACACA | CACCGTGTCCTCATCTTGAAG |
| *Ugt8a* | ACTCCATATTTCATGCTCCTGTG | AGGCCGATGCTAGTGTCTTGA |
| *Plp1* | CCAGAATGTATGGTGTTCTCCC | GGCCCATGAGTTTAAGGACG |
| *Bcas1* | AGAAGCGAAAGGCTCGGAAG | AGGGACAGAATAACTCAGAGTGT |
| *Tet1* | CATTCTCACAAGGACATTCACAACA | AGTAAAACGTAGTCGCCTCTTCCTG |
| *β-actin* | GGCTGTATTCCCCTCCATCG | CCAGTTGGTAACAATGCCATGT |
| *Ccl22* | AGGTCCCTATGGTGCCAATGT | CGGCAGGATTTTGAGGTCCA |
| *Ccl2* | TCAAACTGAAGCTCGCACTCT | GGGGCATTGATTGCATCTGG |
| *Ccr2* | ATCCACGGCATACTATCAACATC | CAAGGCTCACCATCATCGTAG |
| *Cx3cr1* | GAGTATGACGATTCTGCTGAGG | CAGACCGAACGTGAAGACGAG |
| *Itpr2* | CCTCGCCTACCACATCACC | TCACCACTCTCACTATGTCGT |
| *Cacna1a* | CACCGAGTTTGGGAATAACTTCA | ATTGTGCTCCGTGATTTGGAA |
| *Cacna1c* | ATTGTGCTCCGTGATTTGGAA | ACTGACGGTAGAGATGGTTGC |
| *Cacna2d1* | GTCACACTGGATTTTCTCGATGC | GGGTTTCTGAATATCTGGCCTGA |

| | | |
|---|---|---|
| *Cacnb4* | TACCTGCATGGAGTTGAAGACT | TTCGCTCTCTCAAGCTGGATA |
| *Cacng5* | ACCTGGAAGAAGGCATAATCCT | CTATGGTAAAACAGCGTCCTCG |
| *Atp2b1* | AGATGGAGCTATTGAGAATCGCA | CCCTGTAACACGGATTTTTCCTT |
| *Atp2c1* | GCAGGCAGAAGAAGCACCAA | CCTAGTAACCAGCCAACCAAC |
| *Slc8a1* | CTTCCCTGTTTGTGCTCCTGT | AGAAGCCCTTTATGTGGCAGTA |
